# Deep reconstructing generative networks for visualizing dynamic biomolecules inside cells

**DOI:** 10.1101/2023.08.18.553799

**Authors:** Ramya Rangan, Sagar Khavnekar, Adam Lerer, Jake Johnston, Ron Kelley, Martin Obr, Abhay Kotecha, Ellen D. Zhong

## Abstract

Advances in cryo-electron tomography (cryo-ET) have produced new opportunities to visualize the structures of dynamic macromolecular machinery in native cellular environments. Here, we describe a machine learning approach that can reconstruct the structural landscape and dynamics of biomolecular complexes present in cryo-ET subtomograms. This method, cryoDRGN-ET, learns a deep generative model of 3D density maps directly from subtomogram tilt series images and can capture states diverse in both composition and conformation. We use this approach to reconstruct the *in situ* translation dynamics of prokaryotic ribosomes, and we reveal the distribution of functional states during translation elongation populated by *S. cerevisiae* ribosomes inside cells.

## 1 INTRODUCTION

Cryo-electron tomography (cryo-ET) is an imaging technique that provides structural insights spanning cellular to molecular length scales [1, 2]. By computationally combining a series of tilt images of intact cells or thinly milled lamella, cryo-ET can visualize the architecture of whole cells in three dimensions at nanometer resolution. Further computational processing of the resulting 3D tomograms with algorithms for segmentation and subtomogram reconstruction can resolve structures at sub-nanometer resolution, providing detailed snapshots of macromolecular structures and their localization in native contexts [3–8].

A major challenge in image processing workflows for cryo-ET is the analysis of structural heterogeneity within subtomogram data. Subtomogram reconstruction algorithms must cope with imaging attributes specific to cryo-ET such as the extremely low signal-to-noise ratio in exposure-limited individual tilt images, as well as the inherent complexity from variations in conformation and composition of biomolecular complexes within cellular samples taken without purification. While some advanced methods for heterogeneity analysis have been proposed [5, 9–11], the majority of subtomogram processing workflows rely on 3D classification to cluster subtomograms into a few, discrete conformational states. Although this approach has been successfully used to reveal distinct states of macromolecular machines *in situ* [12–14], current processing workflows remain unwieldy, with many manual steps and significant computational requirements. Furthermore, these methods are not well-suited for modeling continuous heterogeneity and require specifying the number of expected states *a priori*, often additionally requiring user-provided masks to focus classification on regions with known variability. More fundamentally, 3D classification requires averaging subtomograms for thousands of particles to obtain well-resolved structures, leading to trade-offs between the number of states that can be reconstructed and the resolution of density maps for these states. While machine learning methods based on deep neural networks have shown recent successes in modeling structural variability in single particle cryo-electron microscopy (cryo-EM) [15–17], their potential has yet to be realized in modeling heterogeneous structures from the cellular milieu.

Here, we introduce cryoDRGN-ET for heterogeneous reconstruction of cryo-ET data (**Fig. 1**). CryoDRGN-ET learns a deep generative model of 3D density maps directly from particle tilt images. Similar to the cryoDRGN method for single particle analysis, cryoDRGN-ET’s generative model is parameterized with a neural field [18] representation of structure that is able to capture diverse sources of heterogeneity, including compositional changes, continuous conformational dynamics, and impurities and artifacts from imaging. Applied to a previously published *in situ* dataset of the bacterial ribosome, cryoDRGN-ET recapitulates the distribution of translational states in quantitative agreement with prior analyses [12], while visualizing continuous motions and membrane-associated conformations in a single model. We then used cryoDRGN-ET to reveal the native structural landscape of the *S. cerevisiae* eukaryotic ribosome from cryo-ET. CryoDRGN-ET is open-source software available in version 3.0 of the cryoDRGN software package (https://cryodrgn.cs.princeton.edu/).

**Fig. 1.**
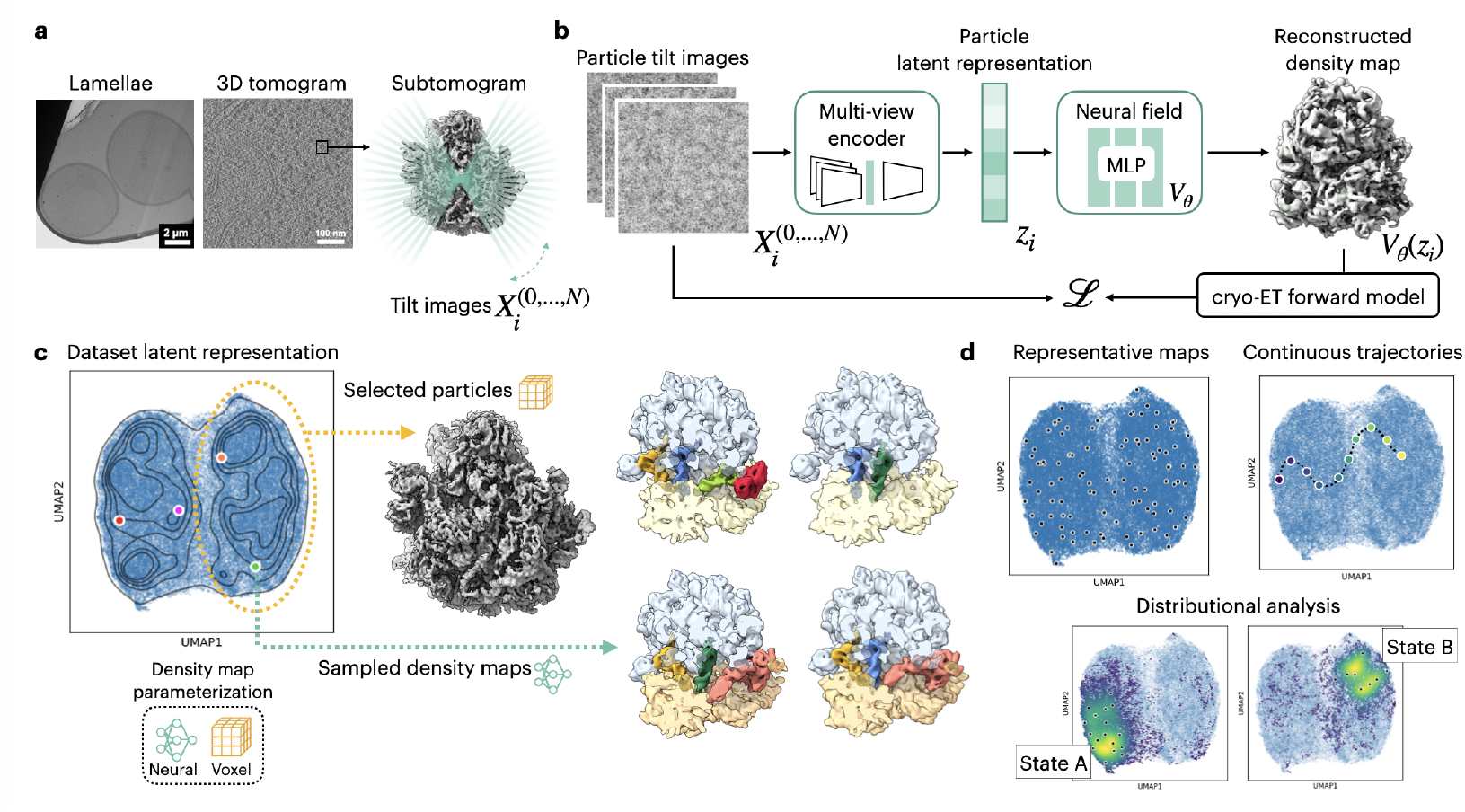
The cryoDRGN-ET method for heterogeneous reconstruction of cryo-ET subtomograms. **a.** Overview of tomography data acquisition and selection of subtomograms, including a schematic showing the series of tilt images that are obtained from each subtomogram. **b.** CryoDRGN-ET architecture. Particle tilt series are transformed into a latent embedding through a multi-view encoder. The decoder includes a multilayer perceptron (MLP) that can reconstruct density maps given a particle’s latent embedding. **c.** Density map generation. Once trained, latent embeddings for a dataset’s particles can be visualized with UMAP and density maps can be generated via two approaches. First, density maps can be directly generated with the parameterized MLP given any latent embedding. For example, four density maps are generated from the colored points in the latent space, with density regions colored to show the ribosome’s large subunit (blue), small subunit (yellow), and different factors binding. Second, density maps can be generated by a standard homogeneous reconstruction from particles selected based on the latent representation. **d.** Analysis of cryoDRGN-ET’s generative model. Density maps can be systematically sampled from the latent representation (here we show k-means clustering centers, k=100), and continuous trajectories between points in latent space can be explored. Density maps of representative states can be classified to further visualize the distributions of particle classes across the latent space.

## 2 RESULTS

### 2.1 Heterogeneous reconstruction of cryo-electron subtomograms with cryoDRGN-ET

CryoDRGN-ET is a generative neural network method for determining a continuous distribution of density maps from cryo-ET subtomogram data. To train cryoDRGN-ET, we use the standard cryo-EM image formation model, extended to apply to tomography (Section 4). Unlike in single particle analysis (SPA) where a single projection image is captured for each particle, in tomography, multiple projections of the same particle are captured from different tilt angles (**Fig. 1a**). CryoDRGN-ET aggregates the different tilt images for each particle with a multiview encoder that outputs a vector *z*_*i*_ ∈ R^*N*^, also referred to as a latent embedding, representing the conformational state of particle *i* (**Fig. 1b**). Then given this latent embedding *z*_*i*_, cryoDRGN-ET’s generative model outputs a 3D density map *V*_*i*_. This map can then be rendered as 2D projections corresponding to each particle’s tilt images given the image pose and estimated CTF parameters. A maximum likelihood objective is used to compare these estimated 2D projections against the observed tilt images. We model high-frequency signal attenuation as a function of electron exposure dose in the CTF for each tilt image [19]. We additionally implement software enhancements for handling large subtomogram datasets containing millions of particle images.

Once training is complete, cryoDRGN-ET provides a per-particle estimate of the dataset’s heterogeneity that can be analyzed through multiple approaches (**Fig. 1c-d**). The distribution of latent embeddings for all particles in the dataset can be visualized in 2D, e.g. with principal component analysis (PCA) or UMAP [20] (**Fig. 1c**). Because cryoDRGN-ET learns a neural representation of 3D density, a representative map can also be generated from any point in the latent space (**Fig. 1c**, Section 4), and these maps can be used to explore and interpret the conformational distribution. For example, to more systematically analyze compositional and conformational heterogeneity, representative density maps can be examined to identify particle classes belonging to distinct states, and continuous trajectories can be generated by sampling maps along paths in the latent space (**Fig. 1d**). Finally, observed states may be validated by selecting constituent particles in particle classes (“distributional analysis”, **Fig. 1d**) and performing a traditional homogeneous reconstruction using voxel-based backprojection, which we newly implement in the cryoDRGN software suite (**Fig. 1c**).

### 2.2 CryoDRGN-ET recapitulates distribution of translation states of the bacterial ribosome

To test cryoDRGN-ET’s subtomogram analysis, we applied it to a previously published *in situ* dataset of the *M. pneumoniae* bacterial ribosome after chloramphenicol (Cm) treatment [3], comparing against prior conventional 3D classification on this dataset [12]. Ribosome particle tilt series images and their associated pose and CTF parameters were obtained with RELION and WARP/M (Section 4). We first assessed these particles’ quality by training cryoDRGN-ET on all 18,466 particles, identifying outliers and particles that produce poor ribosome density maps to exclude in further training runs (**Fig. S1**). We additionally assessed the resolution of the reconstruction when varying the number of tilt images used per particle, finding no further improvement when using more than 8 images per particle (**Fig. S2**). For subsequent cryoDRGN-ET training for this dataset, we used the first 10 tilts in the dose-symmetric tilt acquisition scheme for each particle.

A cryoDRGN-ET network trained on the remaining particles generated density maps displaying known compositional and conformational heterogeneity in the bacterial ribosome, including varying tRNA occupancy at the A-P-E sites, the appearance of elongation factors, subunit rotation, and local motions (**Fig. 2**). Maps from cryoDRGN-ET reproduced the major translational states previously identified in this Cm-treated *M. pneumoniae* ribosome dataset [12]: the P state; EF-Tu-tRNA, P state; A, P state; and A*, P/E state (**Fig. 2a**). These states show density for tRNAs and elongation factor EF-Tu in the expected sites, with significant rotation of the small subunit (SSU) present only in the A*, P/E state as expected [12]. We classified 100 representative density maps across the latent space to assign particles into these 4 states, dividing the latent space into four distinct regions (**Fig. 2b**, Supplementary Video 1, 2). This classification enabled quantifying the relative occupancy of ribosomes in these four states, with most particles representing the A, P state and other minor state populations similar to those found previously by conventional 3D classification [12] (**Fig. 2c**). To validate cryoDRGN-ET density maps and our class assignments, we verified that we could reproduce the structures from homogeneous reconstruction of each state’s particles (**Fig. S3**). In addition, these reconstructions confirmed the rotation of the SSU, with the subunit rotated in the A*, P/E state relative to the A, P state (**Fig. S4**). Since the A, P state included the most particles, a homogeneous reconstruction of this state produced the highest resolution map, with a global estimated resolution of 3.8 Å (**Fig. 2d**).

**Fig. 2.**
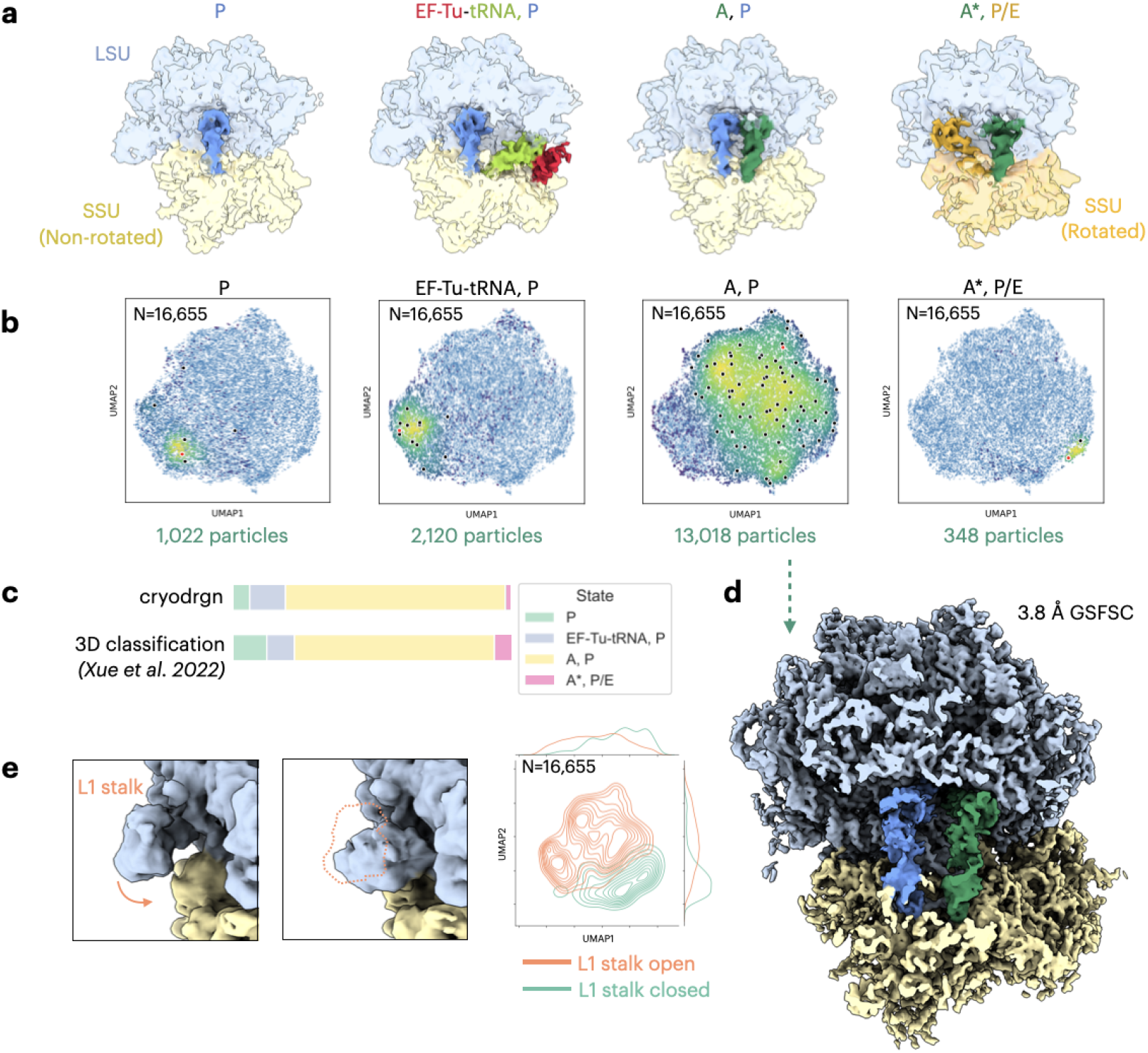
CryoDRGN-ET models translational states from an *in situ* subtomogram dataset of the chloramphenicol-treated *Mycoplasma pneumoniae* 70S ribosome (EMPIAR-10499 [3]). **a.** Representative cryoDRGN-ET maps depicting four translational states. **b.** UMAP visualization of latent embeddings for all particles included in cryoDRGN-ET training, with overlaid heatmaps highlighting the particles belonging to each translational state. Latent embeddings producing representative density maps in **a)** are indicated in red. **c.** Quantification of particle populations in each observed translational state, compared between cryoDRGN-ET and prior conventional 3D classification [12]. **d.** High-resolution reconstruction from voxel-based backprojection in cryoDRGN-ET for particles assigned to the A, P state. **e.** Representative maps from cryoDRGN-ET displaying the L1 stalk open (left) and closed (middle) conformations. A kernel density estimate (KDE) plot displaying the distribution of these two L1 stalk conformations in the latent space (right).

Beyond variation in the tRNA channel and factor-binding sites, cryoDRGN-ET density maps exhibited local dynamics, additional protein factor variability, and larger-scale background variation (Supplementary Video 3). For instance, some cryoDRGN-ET maps showed the L1 stalk in the open state while others included the closed state, with L1 stalk closed conformations overlapping with the A*, P/E state and partially with the A, P state (**Fig. 2e**). We validate the observed L1 stalk conformations with a conventional homogeneous reconstruction from each conformation’s particles (**Fig. S5**). Additionally, some cryoDRGN-ET maps showed density for the N-terminal domain (NTD) of the L7/L12 protein (**Fig. S6a,b**), which is often challenging to resolve on ribosomes with SPA [21]. Finally, cryoDRGN-ET density maps exhibited larger-scale background variation in some cases, showing density for ribosomes bound to the cell membrane in the expected orientation (**Fig. S6a,c**) [22], along with density for neighboring ribosomes in polysomes, also in a canonical orientation (**Fig. S6a,d**) [23]. We found that this large-scale background variation was more visible when using all 41 tilts per particle, as higher tilt angles provide additional views of the variability surrounding each ribosome. We expect that analysis of background variation will be further enhanced when training cryoDRGN-ET on particle sets with larger box sizes that include more surrounding context. Recently, a similar neural network architecture has been proposed to model heterogeneity for tomography [24]. When using this network with default parameters and adding dose exposure settings, we do not observe tRNA or elongation factor heterogeneity in this particle set, with all representative density maps in the A, P state. However, other parameterizations of this network may exhibit more heterogeneity. Finally, unlike the original analysis of this dataset which relied on multiple rounds of 3D classification [12], we note that cryoDRGN-ET is able to recover all states in a single round of training without the use of masks to focus on regions of expected variability.

### 2.3 Revealing the native structural landscape of the *S. cerevisiae* eukaryotic ribosome

We further showcase the method by analyzing a large cryo-ET dataset of the *S. cerevisiae* eukaryotic ribosome collected from lamella that were milled with cryo-plasma focused ion beam milling (cryo-PFIB). With cryoDRGN-ET, we provide the first heterogeneity analysis for the *S. cerevisiae* ribosome from *in situ* cryo-ET, recapitulating known translational states and factor-binding events along with expected continuous conformational motions and spatial background variability. Since we again find that using a subset of tilt images enables high-resolution reconstructions (**Fig. S7a,b**), we carry out all training runs using 10 tilt images per particle for enhanced computational efficiency on this large dataset of 119,031 particles. We begin with a cryoDRGN-ET training run on the complete particle set, finding that the UMAP latent space representation separates particles into three classes, corresponding to rotated SSU, non-rotated SSU, and a group of outlier particles (**Fig 3a**). A homogeneous reconstruction of this outlier group yielded a very noisy map resembling broken particles (**Fig. 3a**), and removing this group of 25,750 particles did not impact the resolution of the consensus refinement (**Fig. S7c**). Homogeneous reconstruction of the two remaining particle classes verified that they corresponded to the SSU rotated and non-rotated states (overlaid in **Fig. 3a**), and a subsequent cryoDRGN-ET training run excluding outlier particles reproduced the separation of SSU rotated and SSU non-rotated particles (**Fig. 3a**). To focus model capacity on further delineating translational states, we trained separate cryoDRGN-ET models on the SSU non-rotated and SSU rotated particles (**Fig. 3b**).

**Fig. 3.**
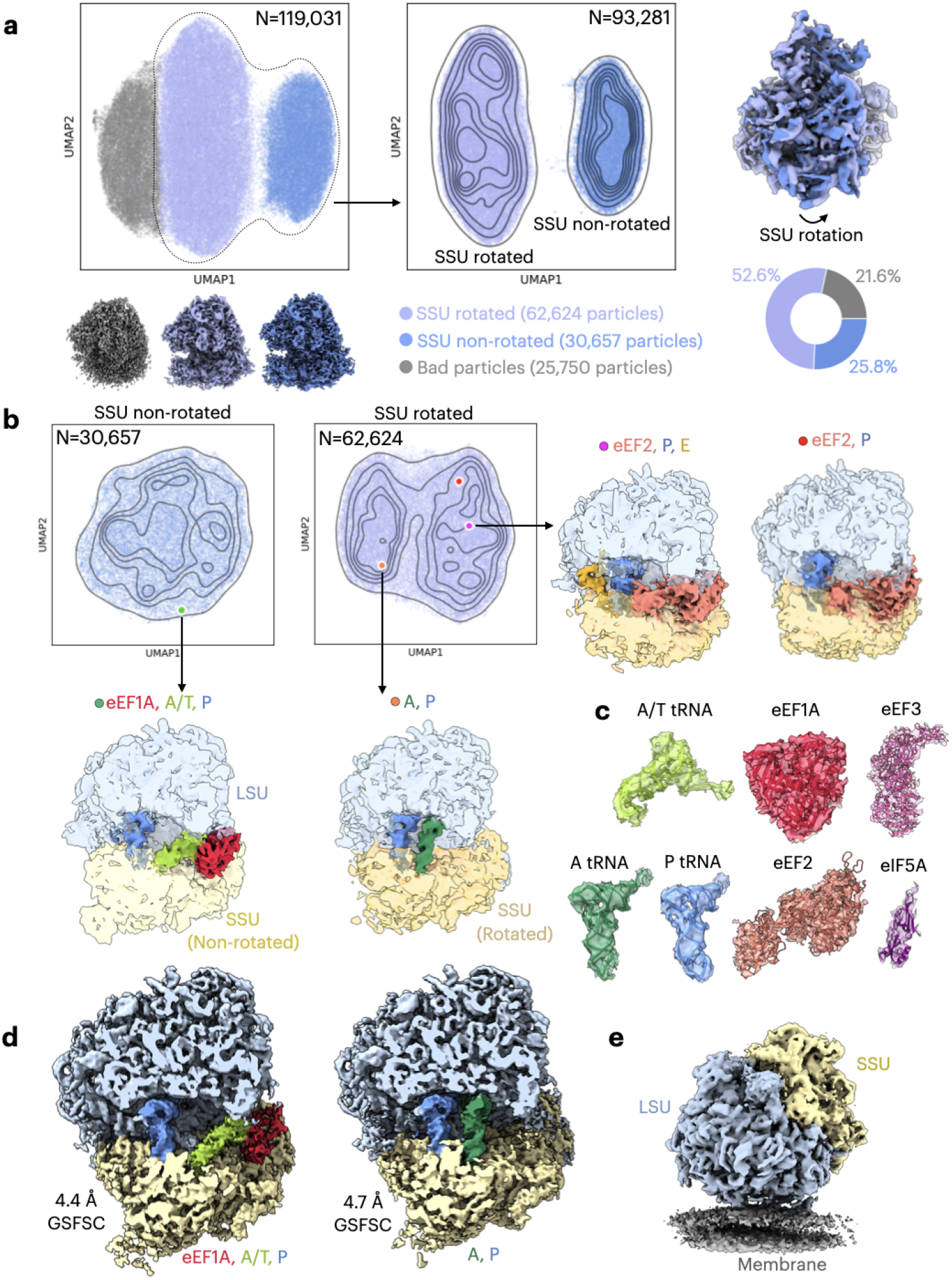
CryoDRGN-ET models *in situ* translation states of the *S. cerevisiae* 80S ribosome. **a.** UMAP visualization of cryoDRGN-ET’s latent space representation of all particles (left) and after excluding a cluster of bad particles (center). Visualizations are shown as scatter plots of particle latent embeddings with the kernel density estimate (KDE) overlaid. Density maps were obtained from a homogeneous reconstruction of particles from the three main clusters (bottom) and overlaid (right) to show the SSU rotation. **b.** UMAP visualization after cryoDRGN-ET training on SSU non-rotated particles (left) and SSU rotated particles (center) with representative cryoDRGN-ET density maps depicting four translational states. Latent embeddings of the representative maps are highlighted in the UMAP visualization. **c.** Atomic models were rigid-body fitted into reconstructed densities from backprojection for the A/T tRNA and eEF1A (in eEF1A, A/T, P state), A and P tRNA (in A, P state), eEF2 (in the eEF2, P, E state), eEF3 (in the eEF1A, A/T, P, eEF3 state) and eIF5A (in the A, P, eIF5A state). **d.** High-resolution backprojection of particles mapped to the eEF1A, A/T, P state (left) and the A, P state (right). **e.** Representative cryoDRGN-ET density map for the membrane-bound ribosome.

Density maps sampled from cryoDRGN-ET training on SSU non-rotated and rotated particles could be classified primarily into four translational states (**Fig. 3b**), providing *in situ* evidence for these functional states and their relative populations in *S. cerevisiae* ribosomes (Supplementary Video 4). Most representative maps from the SSU non-rotated particles corresponded to the eEF1A, A/T, P state, a stage prior to peptidyl transfer. Indeed, the eEF1A, A/T, P state was the most populated across this *S. cerevisiae in situ* ribosome dataset (**Fig. S8a**), agreeing with recent characterization of eukaryotic ribosomes *in situ* from *D. discoideum* and human cells [13, 14]. The conformation of eEF1A and the A/T tRNA in this state is aligned with codon sampling rather than codon recognition (**Fig. S8b**) [14]. Next, we noted a class of representative maps from the SSU rotated particle set corresponding to the A, P state, accounting for the second largest particle population (**Fig. 3b, S9a**). Finally, from the SSU rotated particle set, we noted representative maps corresponding to two post-translocation states: the eEF2, P, E state and the eEF2, P state. We validated particle classification into these four states through homogeneous reconstructions, finding that the resulting reconstruction for each class reproduced expected tRNA and factor density (**Fig. 3c, S8, S9**). When fitting atomic models into these reconstructions, we observed expected SSU motion across these states, with SSU rolling and rotation visible in the A, P state and SSU rotation visible in the post-translocation states (**Fig. S10**). Since the eEF1A, A/T, P state and the A, P state had the highest particle populations, reconstructions from these states produced the highest resolution maps at a global estimated resolution of 4.4 Å and 4.7 Å respectively (**Fig. 3d, S8, S9**).

As in the case of the bacterial ribosome, cryoDRGN-ET was able to further uncover larger-scale background variation and compositional heterogeneity for additional protein factors beyond these canonical translational states (Supplementary Video 5). For instance, representative density maps from cryoDRGN-ET included membrane-bound ribosomes (**Fig. 3e**) and polysomes (**Fig. S11**), with polysome density visible in maps sampled from both SSU rotated and non-rotated states. Moreover, some representative maps included density for the initiation factor eIF5A (**Fig. S12**) [25], and other maps included density for the uL10(P1-P2)_2_ stalk (**Fig. S13**) [26]. Finally, we noted that some representative maps showed the presence of fungal-specific elongation factor eEF3 (**Fig. S14**). eEF3 was present only in representative density maps sampled from SSU non-rotated particles in the eEF1A, A/T, P state, aligning with prior suggestions that eEF3 binding is stabilized in non-rotated states [27]. Density for these additional factors was validated with homogeneous reconstructions of identified particles (**Fig. S12-S14**), for instance with density for eIF5A and eEF3 agreeing with prior atomic models for these factors that were determined from purified samples (**Fig. 3c**). Through these analyses, we show that cryoDRGN-ET was able to model numerous sources of heterogeneity in *S. cerevisiae* ribosomes, providing practical utility with its fast training and interpretable latent space and more fundamentally, a new analytical approach for interrogating structural distributions *in situ*.

## 3 DISCUSSION

In summary, with cryoDRGN-ET, we provide new capabilities for analyzing heterogeneity within cryo-ET subtomograms. This approach leverages the expressive representation power of deep neural networks to generate density maps with compositional and conformational variation from cryo-ET subtomograms. When applied to *in situ* datasets, we characterized translational states of the bacterial ribosome in quantitative agreement with previous results [12], and we newly visualized the native translational states of the *S. cerevisiae* ribosome, confirming the structural characterization from purified systems. Notably, our method yields a distinct estimate of structural heterogeneity for each particle (i.e. *z*_*i*_ or *V*_*i*_) and is able to resolve structural states of interest without requiring any masks designed to focus on expected variability. This relatively unbiased, per-particle heterogeneity estimate can enable the joint analysis of inter- and intra-particle variation, potentially disentangling complex relationships between particles’ conformational states, binding factor composition, and spatial context. Interestingly, we resolve structural heterogeneity using only a small subset of the collected tilt images (i.e. the top 10 highest signal images); a rigorous characterization of the amount of signal needed for representation learning and for high-resolution reconstruction remains a subject of ongoing work. Lastly, our results here relied on image poses obtained from an upstream consensus refinement; the development of cryoDRGN-ET for subtomogram analysis can be coupled with recent developments in neural *ab initio* reconstruction [28, 29], paving the way towards *in silico* purification of dynamic macromolecular machinery within cells.

## 4 ONLINE METHODS

### 4.1 CryoDRGN-ET generative model

CryoDRGN-ET performs heterogeneous reconstruction using a neural network representation for cryo-EM structures. In particular, the central task in cryoDRGN-ET is to learn a function 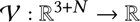 describing an *N* dimensional continuous distribution over 3D cryo-EM density maps. We use a generic latent variable *z* ∈ R^*N*^ to model the conformational distribution and parameterize the generative model with a coordinate-based neural network, *V*_θ_ (γ (k), *z*), where θ are parameters of a multi-layer perceptron (MLP). In cryoDRGN-ET, the density map is specified in the Fourier (or the closely related, Hartley) domain; thus, k ∈ R^3^ are Cartesian coordinates representing Fourier space wavevectors. Similar to recent development in neural fields for modeling 3D signals [18], input coordinates k are expanded in a sinusoidal basis [30]; instead of geometrically-spaced axis-aligned frequencies in cryoDRGN [15, 31], we use frequencies sampled from a Gaussian distribution as in later versions of the cryoDRGN-ET software [32]:

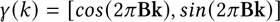

where entries of B ∈ R^*D*×3^ are sampled from N (0, σ^2^) and *D* is a hyperparameter. Without loss of generality, we model density maps from *V*_θ_ on the domain [−0.5, 0.5]^3^ in our coordinate-based neural network. By default, we set σ = 0.5 for our Fourier featurization and set *D* to be the resolution of the training images in pixels.

The generation of cryo-ET subtomogram tilt images follows the standard cryo-EM image formation model with modifications for tomography. We note that many extant methods for *subtomogram averaging* (STA) align and average many subtomogram *volumes* of the same particle. Alternatively, some newer approaches perform STA by directly aligning and averaging the 2D tilt images rather than the subvolumes [3, 4], which avoids artifacts due to the missing Fourier space wedge in individual subtomograms and can be more memory efficient. In cryoDRGN-ET, we treat the subtomograms as cropped 2D tilt series. Thus, the image formation of N tilted images 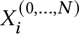 for particle *i* and tilt *j* closely follows that from single particle cryo-EM:

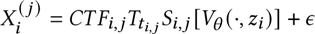

where *CTF_i,j_* applies the contrast transfer function, ε is additive Gaussian white noise, *T_ti,j_* applies a phase shift correponding to translation by *t* ∈ R^2^ in real space, and *S* applies a 2D slicing operator at orientation *R* ∈ *SO*(3) on a volume *V* : R^3^ → R:

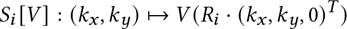

Thus, to generate an image with our coordinate-based neural network, we first obtain oriented 3D coordinates of the 2D central slice corresponding to each pixel from the image, taking a a grid of 3D pixel coordinates originally spanning [−0.5, 0.5]^2^ on the x-y plane and rotating by the pose of each tilt image. Then given these coordinates and the latent embedding predicted for the particle, the volume decoder *V*_θ_ can render a 2D slice in the Fourier (or Hartley) domain. The phase shift corresponding to the 2D real space translation is applied before multiplying by the CTF.

To account for accumulated radiation damage in tomography, we additionally extend the CTF to account for lower signal-to-noise ratio (SNR) in tilts collected at later time-points and higher angles. First, we include a dose exposure correction to account for frequency-dependent signal attenuation in later tilt images, as the sample has been exposed to higher electron doses when these tilts are collected. As described previously [33], for each tilt image we compute this dose exposure correction as 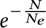, where *N* is the cumulative dose accrued in the sample when this tilt image was collected, and *N*_*e*_ is the dose at which the SNR is 1/*e* of its starting value. Based on previous calibration [33], *N*_*e*_ is computed as 2.81 + 0.245*e*^−1.665*s*^, dependent on spatial frequency *s*. These dose exposure corrections are then masked to 0 for frequencies where the cumulative dose exceeds the frequency-dependent optimal exposure value (2.51284*N*_*e*_) [33]. We multiply the CTF by these dose-exposure corrections during training. Additionally, since sample thickness effectively increases at higher tilt angles leading to decreasing SNR for these tilts, we further multiply the CTF by the cosine of the tilt angle [34]. Our current implementation assumes that data is collected with a dose-symmetric tilt-scheme [35].

### 4.2 CryoDRGN-ET training system

The overall cryoDRGN-ET architecture consists of an image encoder-volume decoder based on the variational autoencoder (VAE) [36]. The above coordinate-based neural network *V*_θ_ serves as the probabilistic decoder. The image encoder, *q*_φ_, embeds cryo-EM image(s) associated with each particle into a lower dimensional latent representation. In cryoDRGN for SPA, an MLP embeds a single image into an *N* dimensional latent vector. In cryoDRGN-ET for tilt series data, the encoder aggregates multiple images of each particle from the tilt series into a single latent vector. The encoder parameterizes a diagonal Gaussian approximate posterior over the latent variable *z*, which we sample from during training, but take the mean value during inference. To embed a series of tilt images, the encoder is split into two MLPs, where the first learns an intermediate embedding of each image, and the second maps the concatenation of the embeddings to the latent space. When experimenting with the number of tilt images that are needed for representation learning and reconstruction, tilt images are ordered by exposure so that the highest signal images are always included.

The training objective is based on the standard VAE objective consisting of a reconstruction error as the squared error between the observed image and a rendered slice from the model and a weighted regularization term on the predicted latent representation as the Kullback-Leibler divergence between the variational posterior and a standard normal prior on *z*. Models are optimized with stochastic gradient descent in minibatches of tilt images from 8 particles using the Adam optimizer [37] with a learning rate of 0.0001. By default, the encoder and decoder MLPs have 3 hidden layers of width 1024 and ReLU activations. For the multiview image encoder, the intermediate embedding dimension for tilt images is 64 by default. We use an 8-dimensional latent variable in all experiments. We use a constant weighting factor *β* of 0.025 on the KL divergence term. For a summary of training and architecture hyperparameters and runtimes in all computational experiments, see Table 1.

**Table 1.** Summary of dataset statistics, training hyperparameters, and runtimes for cryoDRGN-ET training experiments. For neural network architectures *d* × *l*, *d* indicates the number of nodes per layer and *l* indicates the number of hidden layers. Training times were recorded on the indicated number of A100 GPUs.

| Dataset | Particle class | # Particles | # Tilts per particle | \|z\| | Encoder architecture | Decoder architecture | Epochs | Training time | # GPUs |
| --- | --- | --- | --- | --- | --- | --- | --- | --- | --- |
| *M. pneumoniae* | All | 18,466 | 1 | 8 | 256x3 | 256x3 | 50 | 13 min | 4 |
| *M. pneumoniae* | Filtered | 16,655 | 10 | 8 | 1024x3 | 1024x3 | 50 | 3 hours 35 min | 1 |
| *M. pneumoniae* | Filtered | 16,655 | 41 | 8 | 1024x3 | 1024x3 | 50 | 12 hours 55 min | 1 |
| *S. cerevisiae* | All | 119,031 | 10 | 8 | 1024x3 | 1024x3 | 50 | 20 hours 32 min | 4 |
| *S. cerevisiae* | Filtered | 93,281 | 10 | 8 | 1024x3 | 1024x3 | 50 | 18 hours 36 min | 1 |
| *S. cerevisiae* | SSU non-rotated | 30,657 | 10 | 8 | 1024x3 | 1024x3 | 50 | 6 hours 8 min | 1 |
| *S. cerevisiae* | SSU rotated | 62,624 | 10 | 8 | 1024x3 | 1024x3 | 50 | 12 hours 12 min | 1 |

### 4.3 Voxel-based homogeneous reconstruction

To enable validation of particle selections, we implemented conventional voxel-based backprojection to reconstruct density maps given particles’ tilt images and poses. We populate volume slices in Fourier space with the Fourier transform of each image based on its pose, applying the CTF using the Wiener filter. The 3D reconstruction is computed from the backprojected slices as previously described in [33, 38] as

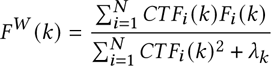

The optimal value of *λ*_k_ is 1/*SNR(k)*, which can be estimated from the data [38]. However, we found that this led to over-regularization in the absence of solvent masking, and we achieved acceptable results with a constant regularization across frequencies equal to the average of the unregularized denominator across voxels

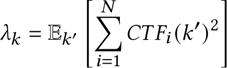

As with cryoDRGN-ET training, we apply dose-exposure and tilt angle corrections to the CTF when carrying out voxel-based backprojection.

To calculate gold-standard Fourier shell correlation (GSFSC) curves [39], we use a custom script implemented in the cryoDRGN-ET software, with backprojected maps from two random halves of the dataset. Prior to calculating FSC curves, we apply soft real-space masks that were obtained as previously described [15]. In particular, masks are defined by first thresholding the full dataset’s consensus density map at half of the 99.99th percentile density value. The mask is then dilated by 25 Å from the original boundary, and a soft cosine edge is used to taper the mask to 0 at 15 Å from the dilated boundary.

### 4.4 Bacterial ribosome dataset pre-processing (EMPIAR-10499)

Raw tilt movies were processed in Warp [3], where motion correction and patch CTF estimation were performed. The tilt-series stack was generated from Warp and the tilt-series were aligned using AreTomo [40]. The tilt-series CTFs were estimated in Warp and tomograms were reconstructed in Warp at a pixel size of 10 Å, where the tomograms were denoised to enhance the contrast for particle picking. Nine denoised tomograms were manually picked in crYOLO and used to train a crYOLO model [41]. In total, an initial 32,253 particle locations were found, and the subtomograms were extracted at a pixel size 10 Å with a box size of 64 pixels. Approximately 500 subtomograms were extracted at 10 Å, and an initial model was generated using the VDAM algorithm in RELION 4.0 [4]. Multiple rounds of 3D classification were performed using the generated initial model to remove obvious bad particles, filtering the dataset to 25,102 particles. These subtomograms were then extracted in Warp at a pixel size of 5 Å with a box size of 128 pixels. One more round of 3D classification was performed, where 18,326 subtomograms were selected and subjected to an initial alignment in RELION 4.0 3D-autorefine with a mask surrounding the large and small subunit. These subtomograms were then extracted in Warp at 1.705 Å, with a box size of 294, where multi-particle refinement was performed in M [3] with a binary mask encompassing the large and small subunit of the ribosome. Global movement and rotation with a 5 × 5 × 41 image-space warping grid, a 8 × 8 × 2 × 10 volume-space warping grid and particle pose trajectories with three temporal sampling points were refined with 5 iterations. Starting at the third iteration, CTF parameters were also refined, and at iteration 4, reference-based tilt-movie alignment was performed in M. This resulted in a 3.6 Å reconstruction of the *Mycoplasma pneumoniae* 70S ribosome.

### 4.5 Bacterial ribosome cryoDRGN-ET analysis

#### 4.5.1 Particle filtering

In the initial analysis of this dataset, a standard single-particle cryoDRGN model (software version 2.3.0) was trained on the 18,655 0-degree tilt images (D=128, 3.9 Å/pix) [15]. The encoder and decoder architectures had 3 hidden layers of width 256 (denoted 256 × 3), and the latent variable dimension was 8. The model was trained for 50 epochs across 4 A100 GPUs, taking 13 minutes total. Once trained, cryoDRGN’s analysis pipeline (“cryodrgn analyze”) was used to visualize the latent space and produce representative density maps. Outliers were removed using cryoDRGN’s interactive lasso-tool on the UMAP visualization of the latent embeddings, leading to a filtered dataset of 16,655 particles. A consensus refinement of the filtered dataset yielded the same global resolution map.

#### 4.5.2 Reconstruction with varying number of tilts

We carried out separate voxel-based backprojections for the filtered dataset of 16,655 particles when using 1, 2, 5, 8, 10, 16, 32, and 41 tilts per particle. When using a subset of tilts, tilts were chosen to be those with lowest dose exposure (collected earliest in the tilt-series.) Local resolution estimates were made in RELION 4.0 [42].

#### 4.5.3 cryoDRGN-ET training

A multi-tilt cryoDRGN-ET model was trained on the filtered dataset of 16,655 particles for 50 epochs taking 3 hours and 35 minutes on 1 GPU, with the top 10 tilts used during training (D=128, 3.9 Å/pix). The encoder and decoder architectures were 1024 × 3, and the latent variable dimension was 8. We additionally trained a cryoDRGN-ET model with all 41 tilts per particle used during training (D=128, 3.9 Å/pix) taking 12 hours and 55 minutes on 1 GPU, using the same filtered particle set and architecture settings.

#### 4.5.4 Classification

After cryoDRGN-ET training, the distribution of structures from each training run was systematically sampled by using the “cryodrgn analyze” pipeline with *k* = 100, where 100 density maps are generated at k-means cluster centers of the latent embeddings as described previously [15]. The 100 density maps from the training run with 10 tilts per particle were then manually classified into 4 states based on tRNA and elongation factor occupancy. Additionally, all 100 density maps were classified into either L1 open or L1 closed conformations. A representative structure of each state was manually selected for visualization in **Fig. 2**. Additional representative density maps with membrane-bound ribosomes, polysomes, and the NTD of L7/L12 visible were selected from the 100 representative density maps for the 41 tilt training run (**Fig. S6**).

#### 4.5.5 High resolution reconstruction and validation

To validate each state, the particles corresponding to each selected cluster from k-means clustering were combined. We then backprojected the tilt images from the high-resolution dataset (D=294, 1.7 Å/pix) using “cryodrgn backproject_voxel”. GSFSC curves between half-maps were obtained as described above to assess resolution. High-resolution backprojections were low-pass filtered to the resolution from GSFSC curves for visualization.

#### 4.5.6 Visualization

To visualize factors bound in the density map reconstructions and selected representative structures from cryoDRGN-ET (**Fig. 2a,d**), we dock in atomic models that were previously determined based on density maps from conventional 3D classification of this dataset in RELION [12]. For the A, P state we used PDB ID 7PHB; for the EF-Tu-tRNA, P state we used PDB ID 7PHA; for the A*, P/E state we used PDB ID 7PHC; and for the P state we used PDB ID 7PH9 [12]. Models were fit into maps and colored by zone in ChimeraX [43].

### 4.6 *S. cerevisiae* sample preparation

*S. cerevisiae* cells were grown in log phase conditions to an OD600 of 0.8. 4 µL of the cells were applied to a glow-discharged 200 mesh holey carbon grid copper grid (Quantifoil R1.2/3) and vitrified in a liquid ethane using Vitrobot Mark IV (Thermo Scientific) set at 4 °C and 100% humidity. Settings: blot force = 10; blot time = 10 s; wait time = 1 s. Samples were stored under liquid nitrogen until use. Grids were clipped in slotted Autogrids (Thermo Fisher Scientific) and subjected to automated lamella preparation using an Arctis cryo plasma FIB (Thermo Fisher Scientific) with AutoTEM Cryo software (Thermo Fisher Scientific) as described elsewhere. Prior to milling, grids were coated with a layer of ion-sputtered, metallic platinum (Pt) for 30 s (Xe+, 12 kV, 70 nA). This was followed by 400 nm cry-deposition of organometallic Pt using the gas injection system, then an additional ion-sputtered platinum layer (Xe+, 12 kV, 70 nA, 120 sec). Next, grids were surveyed using Maps software (Thermo Fisher Scientific) for lamella site identification followed by automated lamella preparation using AutoTEM Cryo with a final thickness range set between 100-250nm. All FIB milling were performed using xenon. After the final milling step, the lamellae were again sputter coated with a thin layer of ion-sputtered metallic Pt (Xe+, 12 kV, 30 nA, 8 sec).

### 4.7 *S. cerevisiae* dataset acquisition

Datasets were collected using a Krios G4 equipped with XFEG, Selectris X energy filter and Falcon 4 direct electron detector (Thermo Fisher Scientific). Tilt-series were collected with a dose-symmetric tilt scheme [35] using TEM Tomography 5 software (Thermo Fisher Scientific). The tilt span of ± 60° was used with 3° steps, starting at ± 10° to compensate for the lamella pre-tilt. Target focus was changed for each tilt-series in steps of 0.25 µm over a range of −1.5 µm to −3.5 µm. Data were acquired in EER mode of Falcon 4 with a calibrated physical pixel size of 1.96 Å and a total dose of 3.5 e-/Å^2^ per tilt over ten frames. A 10 eV slit was used for the entire data collection. Eucentric height estimation was performed once for each lamella using stage tilt method in TEM Tomography 5 software. Regions of interest were added manually, and positions saved. Tracking and focusing were applied before and after acquisition of each tilt step. The energy filter zero-loss peak was tuned only once before starting the data acquisition.

### 4.8 *S. cerevisiae* dataset pre-processing

The data were preprocessed using TOMOgram MANager (TOMOMAN) [44] and the following external packages were used therein. EER images were motion corrected using a modified implementation of RELION’s motioncor [45]. The defocus was estimated using tiltCTF as implemented within TOMOMAN (tiltCTF uses CTFFIND4 [46] for some steps). Tiltseries were aligned using fiducial-less alignment in AreTomo [40]. Initial tomograms without CTF correction were reconstructed using IMOD’s tilt package [47]. 3D CTF corrected tomograms were reconstructed using novaCTF [48] at 8x binning and used for template matching.

Initial particle positions for 80S ribosomes were determined using the noise correlation template matching approach implemented in STOPGAP [7]. PDB entry 6GQV [49] for 80S ribosomes was used to generate a template using the simulate command in cisTEM [50]. Approximately 1000 particles per tomogram were picked from 260 tilt series. Subsequent subtomogram averaging and classification were performed using STOPGAP. 3D classification was performed using simulated annealing stochastic hill climbing multi-reference alignment as described before [11]. The resulting 130K particles were then exported to Warp [51] using TOMOMAN [44]. Subtomograms were reconstructed in RELION 3.0 [52] using Warp at 2x binning (3.92 Å/pix). An iterative approach with subtomogram alignment and additional 3D classification in RELION and tilt-series refinement in M was performed until no further improvement in the gold standard Fourier Shell Correlation (GSFSC) resolution or the map quality was observed. For final averages, 119k particles were reconstructed at an unbinned pixel size of 1.96 Å, and another round of subtomogram alignment in RELION and tilt-series refinement in M [3] were performed until no further improvement in GSFSC resolution or the map quality was observed. For the LSU focused reconstruction and the subsequent analysis of the structural heterogeneity of the 80S ribosome, an additional round of subtomogram alignment in RELION and subsequent tilt-series refinement in M were performed using a focused mask around LSU. The final set of 119k particles were then extracted as 2D sub-tilt-series at binning 1x and 2x using Warp, and used for analyzing conformational heterogeneity with cryoDRGN-ET.

### 4.9 *S. cerevisiae* ribosome cryoDRGN-ET analysis

#### 4.9.1 CryoDRGN-ET training: full dataset

A cryoDRGN-ET model was trained on the full dataset of 119,031 particles for 50 epochs, with the top 10 tilts used during training (D=128, 3.92 Å/pix). The architectures of the two encoder MLPs and decoder MLP were 1024 × 3, and the latent variable dimension was 8. The model was trained for 50 epochs across 4 A100 GPUs, taking 20 hours and 32 minutes total. Once trained, cryoDRGN-ET’s analysis pipeline (“cryodrgn analyze”) was used to visualize the latent space and produce representative density map. We sampled both 20 structures for initial visualization and 100 density maps for a more comprehensive assessment. The UMAP visualization of the latent space revealed three clusters of particles which were assigned as 1) outliers, 2) the SSU rotated particles and 3) the SSU non-rotated particles by visual inspection of representative density maps from each cluster. Particles corresponding to each cluster were selected using cryoDRGN-ET’s interactive lasso-tool on the UMAP visualization of the latent embeddings. A homogeneous reconstruction of each set of particles was then performed with “cryodrgn backproject_voxel” (**Fig. 3a**).

#### 4.9.2 Reconstruction experiments

We carried out voxel-based backprojections for the dataset of 93,281 SSU rotated and non-rotated particles when using 1, 2, 5, 8, 10, 16, and 32 tilts per particle. We did not explore using all 41 tilts for these comparisons and further experiments on this dataset, as many particles did not have all 41 tilt images available. We additionally carried out voxel-based backprojections with all available tilts per particle for both the full dataset of 119,031 particles and the filtered set with 93,281 particles to assess the effects of particle filtering. As before, when using a subset of tilts, tilts were chosen to be those with lowest dose exposure (collected earliest in the tilt-series.) Local resolution estimates were made in RELION 4.0 [42].

#### 4.9.3 CryoDRGN-ET training: hierarchical analysis

Three additional cryoDRGN-ET models were trained on the remaining good particles (93,281 particles) (**Fig. 3a**), the SSU rotated state (62,624 particles) and SSU non-rotated state (30,657 particles) (**Fig. 3b**). All training runs were carried out for 50 epochs, with latent variable dimension 8 and encoder and decoder MLP dimensions of 1024 × 3. The training run on all SSU rotated and non-rotated particles took 18 hours and 36 minutes on 1 A100 GPU, the training run on the SSU rotated particles alone took 12 hours and 12 minutes on 1 A100 GPU, and the training run on the SSU non-rotated particles alone took 6 hours and 8 minutes on 1 A100 GPU.

#### 4.9.4 Classification

After cryoDRGN-ET training, the distribution of structures from each training run was systematically sampled by using the “cryodrgn analyze” pipeline with *k* = 100, where 100 density maps are generated at k-means cluster centers of the latent embeddings. For the two training runs that separately processed SSU rotated particles and SSU non-rotated particles, we classified all 100 representative density maps into corresponding translational states. To classify density maps, we docked in ribosome structures (PDB IDs: 3J7R [53], 5LZS [25], 6GQV [49], 6TNU [54]) that included the following tRNA and elongation factors: A tRNA, P tRNA, E tRNA, A/P tRNA, P/E tRNA, A/T tRNA, eEF2, eEF1A. We then inspected maps to identify the presence of factors. A representative structure for each state was manually selected for visualization in **Fig. 3**. We additionally identified all representative density maps that included density for eIF5A, for eEF3, and for uL10 and the NTD of P1 and P2. From both of these training runs, we further identified representative density maps that included partial density for polysomes. Finally, from the training run that included both SSU rotated and non-rotated particles together, we identified a membrane-bound representative ribosome map.

#### 4.9.5 High resolution reconstruction and validation

To validate each state, the particles corresponding to each selected cluster center from k-means clustering were combined. We then backprojected the tilt images from the high-resolution dataset (D=256, 1.96 Å/pix) using “cryodrgn backproject_voxel”. We compute gold-standard Fourier shell correlation (GSFSC) curves [39] between half-maps to assess resolution. High-resolution backprojections were low-pass filtered to the GSFSC_0_.143 resolution for visualization.

#### 4.9.6 Visualization

To color factors in representative density maps and reconstructions (**Fig. 3**b), and to visualize the fit of individual factors in density (**Fig. 3**c), atomic models for each state were assembled by docking into high-resolution reconstructions. For each translational state, we obtained atomic models for the elongation factors and tRNAs separately, along with separate atomic models for the LSU and SSU, and we docked each of these models as rigid bodies into the high-resolution reconstruction density maps with ChimeraX [43]. For the eEF1A, A/T, P state, we obtained atomic models for eEF1A, the A/T tRNA, and the P tRNA from PDB ID: 5LZS [25] and the large subunit (LSU) and small subunit (SSU) from PDB ID: 3J78 [55]. For the A, P state, the A tRNA, P tRNA, LSU, and SSU were obtained from PDB ID: 6TNU [54]. For the post-translocation states, the eEF2, E tRNA, LSU, and SSU were obtained from PDB ID: 6GQV [49], and the P tRNA was obained from 6TNU [54]. For fitting in an atomic model for eIF5A, we used eIF5A model from PDB ID: 5LZS [25]. For fitting in an atomic model for eEF3, we used the eEF3 model from PDB ID: 7B7D [27].

To observe SSU rotation in (**Fig. S10**), we fit atomic models for the SSU head, SSU body, and LSU separately into high-resolution reconstructions for each visualized translation state. These three rigid bodies were obtained from an atomic model of the *S. cerevisiae* ribosome with the SSU in a non-rotated state (PDB ID: 3J78 [55]). After these three rigid bodies were sequentially fit into each density map in ChimeraX [43], the resulting atomic models were aligned with PDB ID: 3J78 on the large subunit to visualize SSU rotation and rolling. To visualize the SSU head swivel, models containing the SSU head and body alone were aligned on the SSU body.

## 5 DATA AVAILABILITY

All *S. cerevisiae* raw data will be deposited to EMPIAR. Maps will be deposited to EMDB and cryoDRGN model weights and any associated files needed to reproduce this analysis will be deposited to Zenodo. Atomic models used from previous studies were obtained from the PDB (7PHA, 7PHB, 7PHC, 7PH9, 3J7R, 5LZS, 6GQV, 6TNU, 3J78, and 7B7D).

## 6 CODE AVAILABILITY

Software is available at https://github.com/ml_struct_bio/cryodrgn in version 3.0.0-beta.

## 7 ACKNOWLEDGEMENTS

We thank Vineet Bansal and Michal Grzadkowski for software engineering support, Ryan Feathers for assistance with the manuscript, Niels Fischer for feedback and insight on ribosome structures, Fred Hughson for feedback on the manuscript, computational resources and support from Princeton Research Computing, and Princeton University startup funds for support of this work.

## 8 AUTHOR CONTRIBUTIONS

AK and EZ conceived of the work and supervised; RR, AL, and EZ implemented the methods and performed the computational experiments; RK and AK prepared yeast samples and collected cryo-ET data; SK, JJ, and MO processed data; RR and EZ wrote the manuscript with feedback from all authors.

## 9 COMPETING INTERESTS STATEMENT

RK, MO, and AK are employees of Thermo Fisher Scientific, a commercial entity that sells instrumentation used in this study.

## A SUPPLEMENTARY FIGURES

**Fig. S1.**
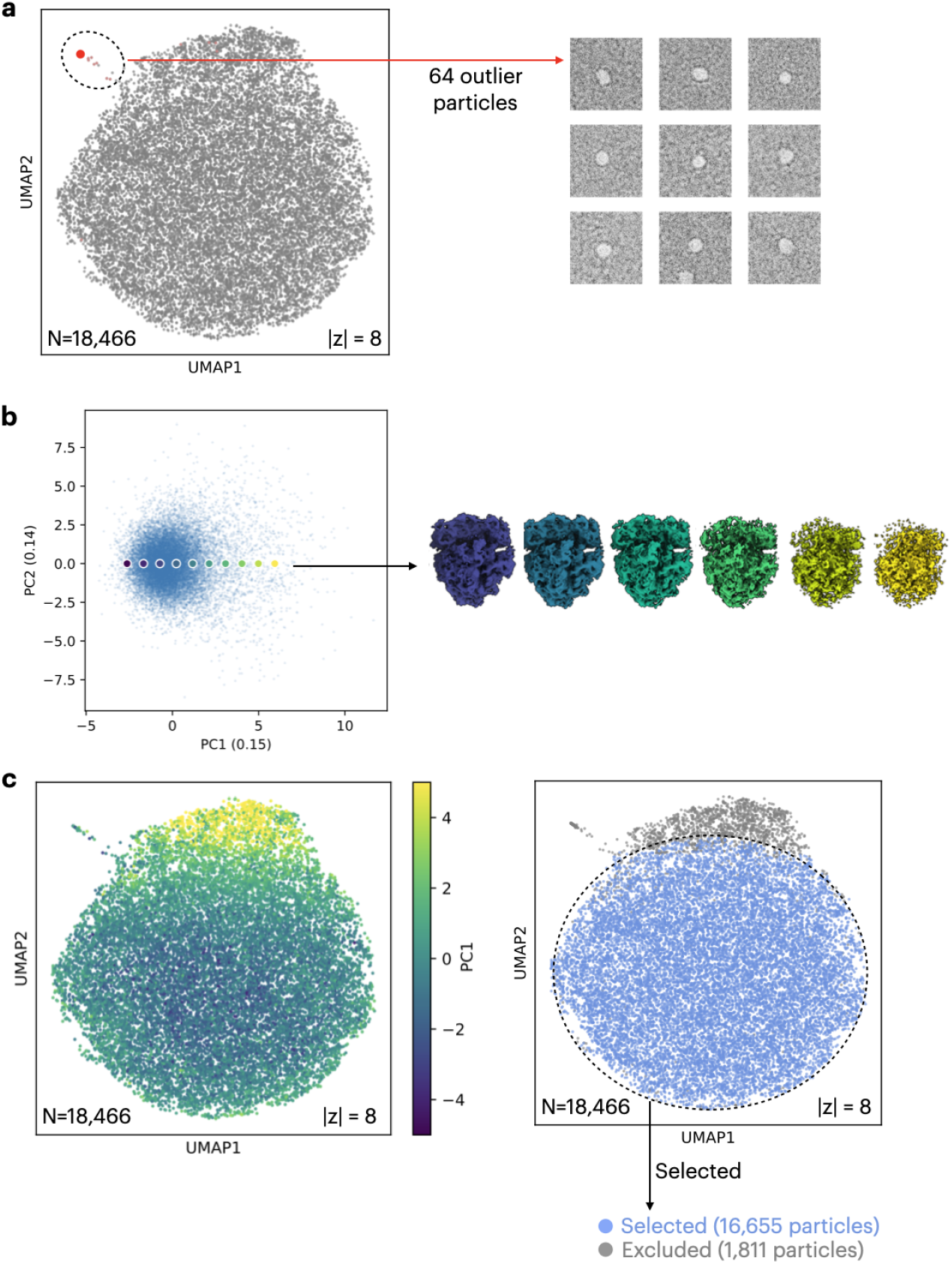
Latent space-based filtering of the *M. pneumoniae* bacterial ribosome subtomograms. **a.** UMAP visualization of cryoDRGN’s latent space representation from a training run using all particles (D=128, 3.9 Å/pixel), showing example particle images for the highlighted group of outlier particles. **b.** Visualization of the latent space along the first and second principal components (PCs), showing representative density maps for the highlighted traversal across PC1. **c.** UMAP visualization of the latent space for the same cryoDRGN training run colored by PC1 using the same coloring as in **b)** (left), and colored by a particle selection that excludes outlier particles (right).

**Fig. S2.**
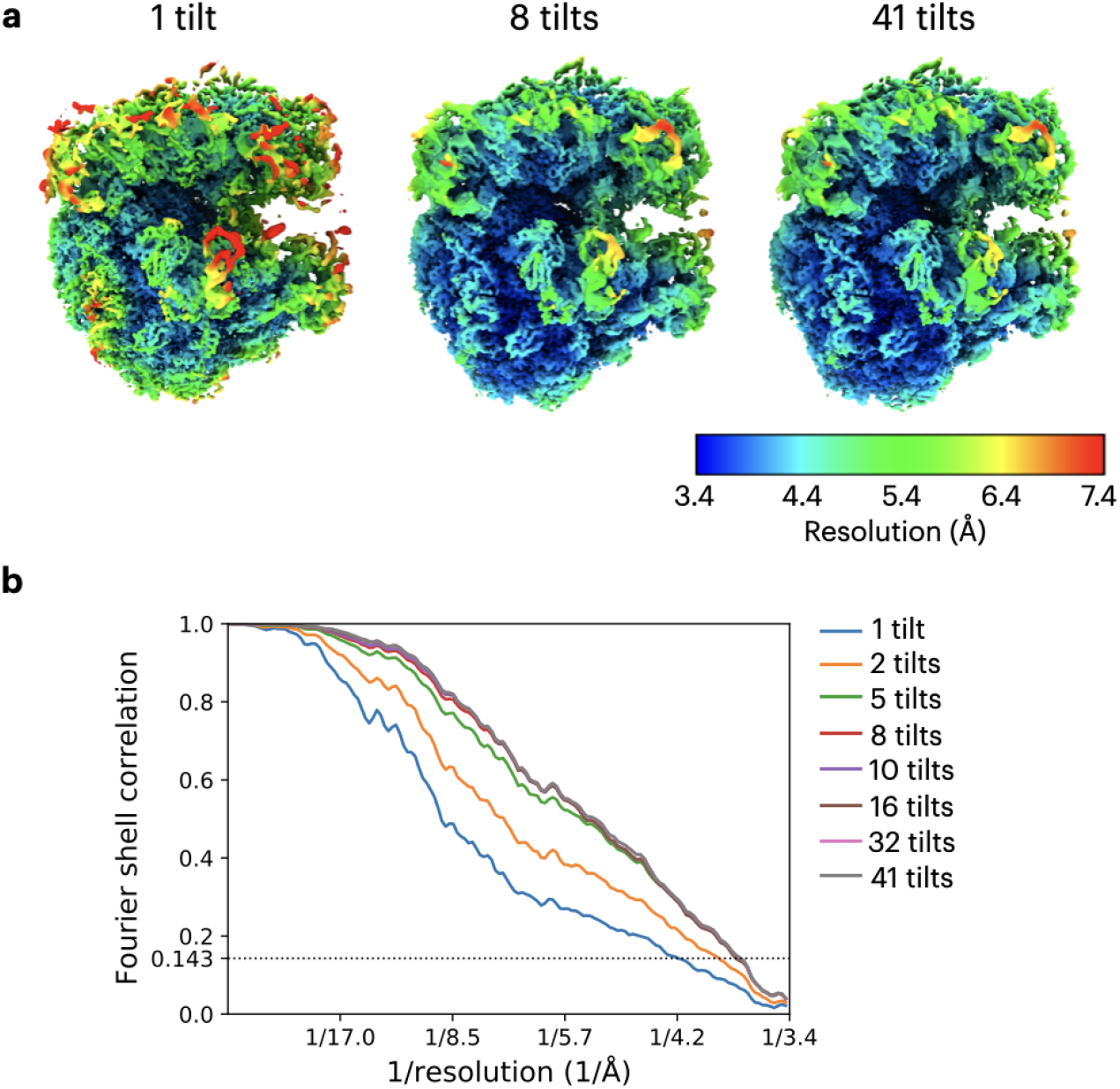
Homogeneous reconstruction of the *M. pneumoniae* ribosome varying the number of tilt images (D=294, 1.7 Å/pixel). **a.** Local resolution estimated from RELION 4.0 [42] for reconstructions using 1 tilt, 8 tilt, and 41 tilts per particle, with maps obtained through voxel-based backprojection in cryoDRGN-ET. **b.** GSFSC curves for varying numbers of tilts per particle.

**Fig. S3.**
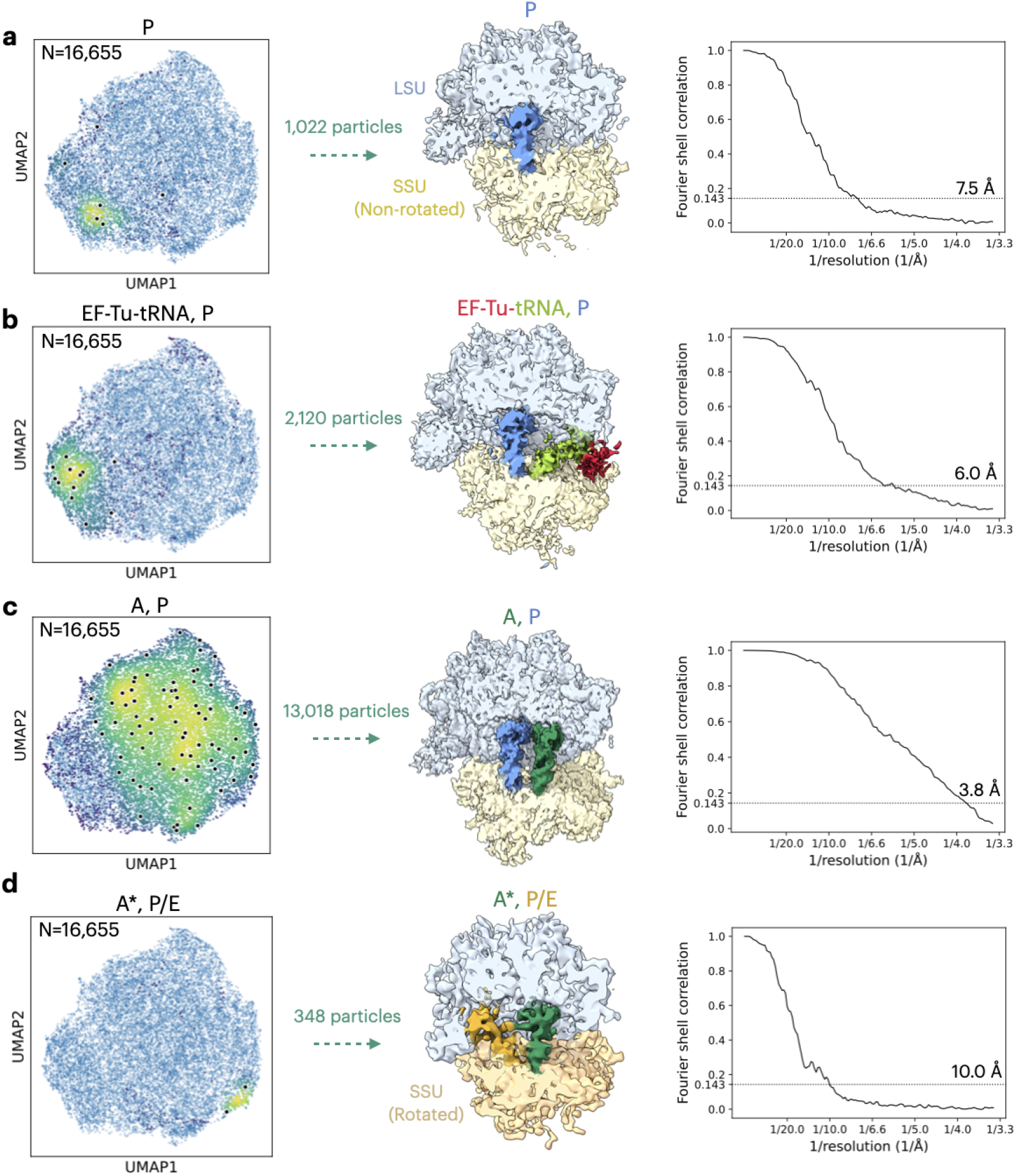
Homogeneous reconstruction of translational states of the *M. pneumoniae* ribosome identified in cryoDRGN-ET, in the following states: **a.** the P state; **b.** the EF-Tu-tRNA, P state; **c.** the A, P state; and **d.** the A*, P/E state. For each state, the left panel shows the UMAP visualization of the latent space from cryoDRGN-ET training of the filtered particle set for the *M. pneumoniae* dataset, with overlaid heatmaps highlighting particles belonging to each state. The middle column depicts the homogeneous reconstruction (D=294, 1.7 Å/pixel) for particles selected in each state using cryoDRGN-ET’s voxel-based backprojection. Reconstructions are low-pass filtered to the GSFSC resolution (0.143 cutoff) and colored by corresponding factors. In the right column, GSFSC curves are depicted for each case.

**Fig. S4.**
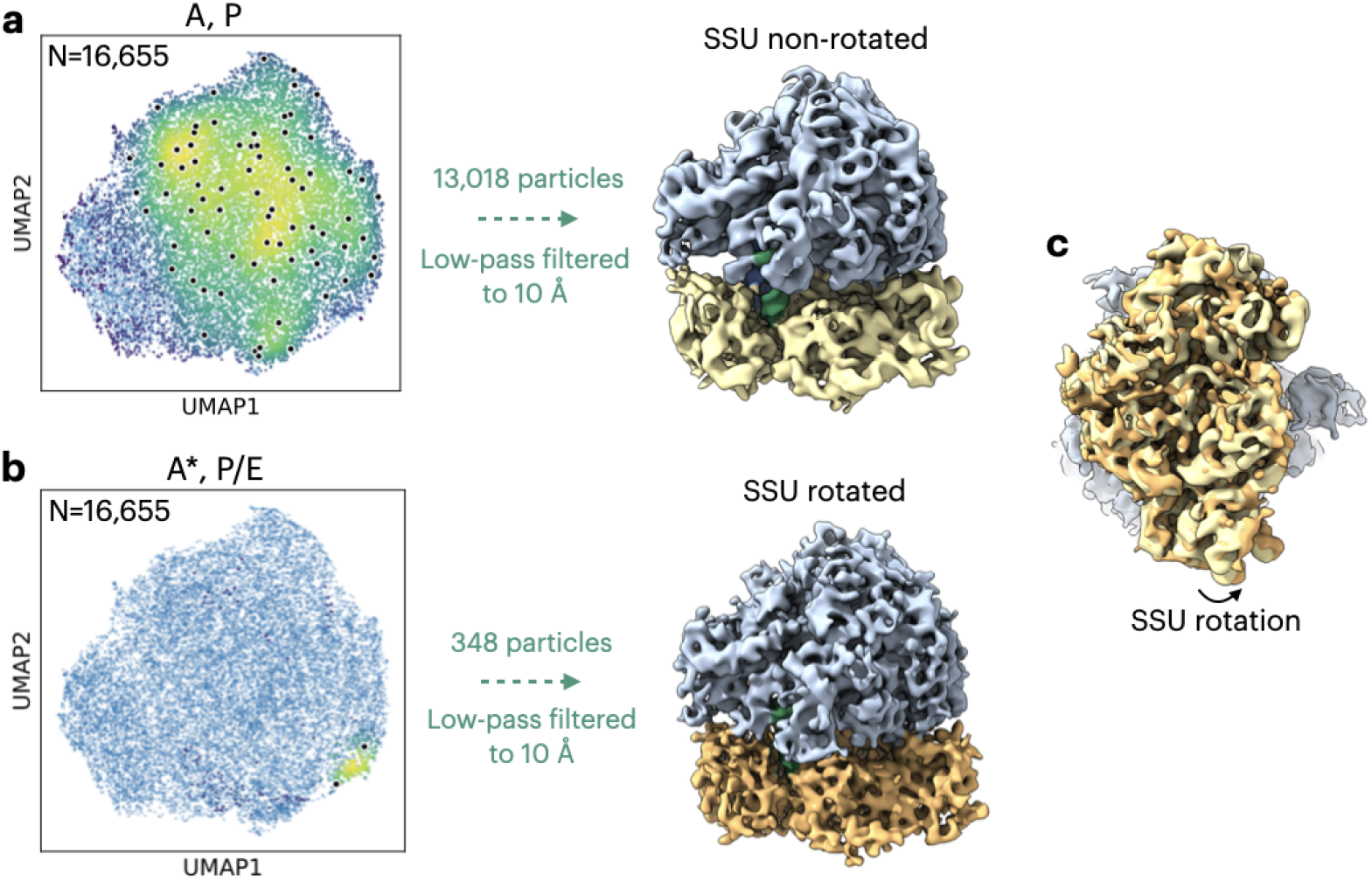
SSU rotation of the *M. pneumoniae* ribosome identified in cryoDRGN-ET. In the left column is the UMAP visualization of the latent space from cryoDRGN-ET training on the filtered particle set, with a heatmap overlaid depicting the distribution of particles in **a.** the A, P state, and **b.** the A*, P/E state. In the middle column are density maps obtained by voxel-based backprojection of particles (D=294, 1.7 Å/pixel) from these two states low-pass filtered to 10 Å resolution. In the right column, these two reconstructions are overlaid and viewed facing the SSU to depict SSU rotation.

**Fig. S5.**
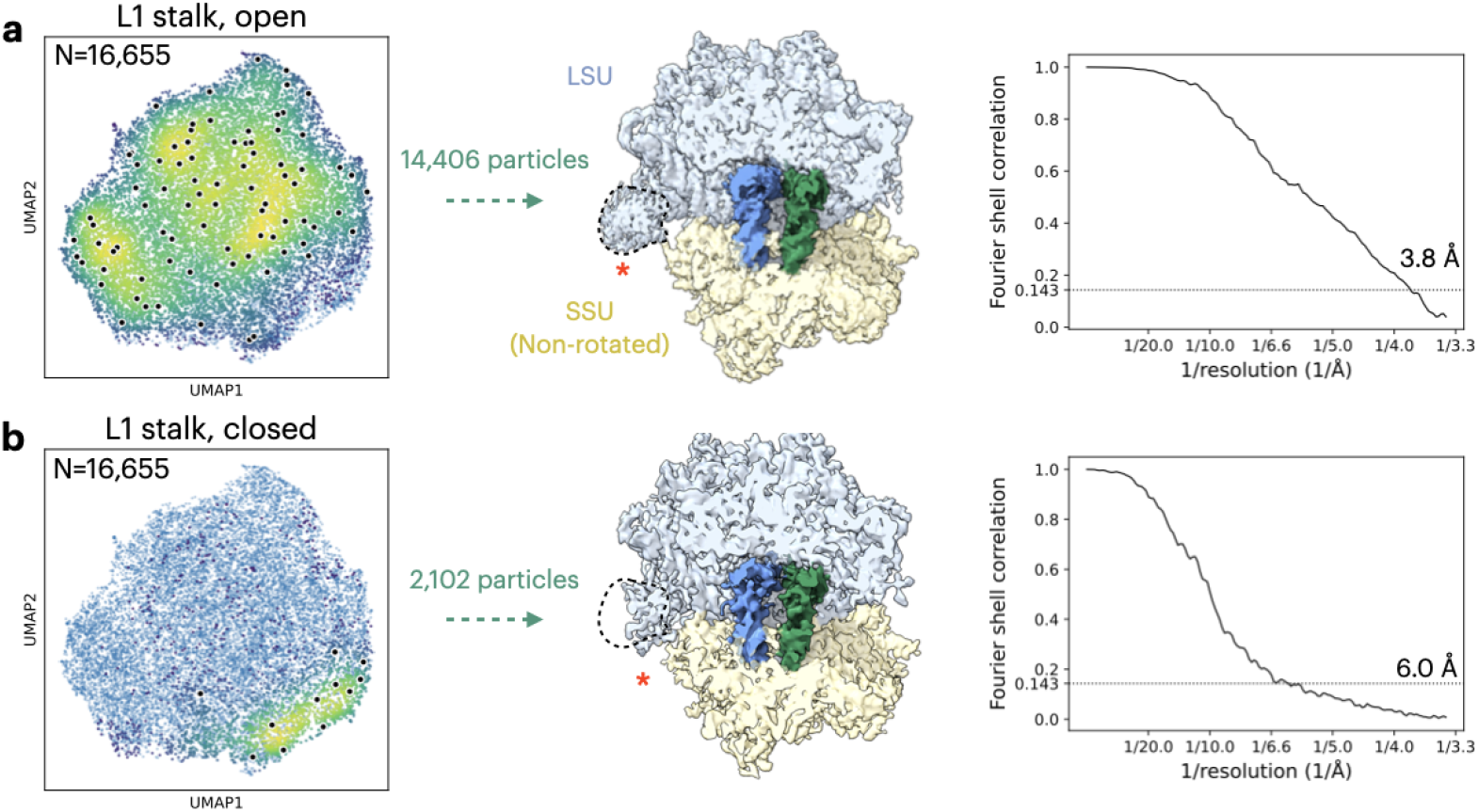
L1 stalk motion of the *M. pneumoniae* ribosome identified in cryoDRGN-ET. In the left column is the UMAP visualization of the latent space from cryoDRGN-ET training on the filtered particle set, with a heatmap overlaid depicting the distribution of particles in **a.** the L1 stalk open conformation and **b.** the L1 stalk closed conformation. In the middle column are high-resolution reconstructions (D=294, 1.7 Å/pixel) obtained by voxel-based backprojection of particles from these two states low-pass filtered to the GSFSC resolution. The L1 stalk in both density maps is highlighted with a red asterisk, and a dotted line indicates the position of the open L1 stalk conformation overlaid on the closed L1 stalk reconstruction. In the right column are GSFSC curves for these two states.

**Fig. S6.**
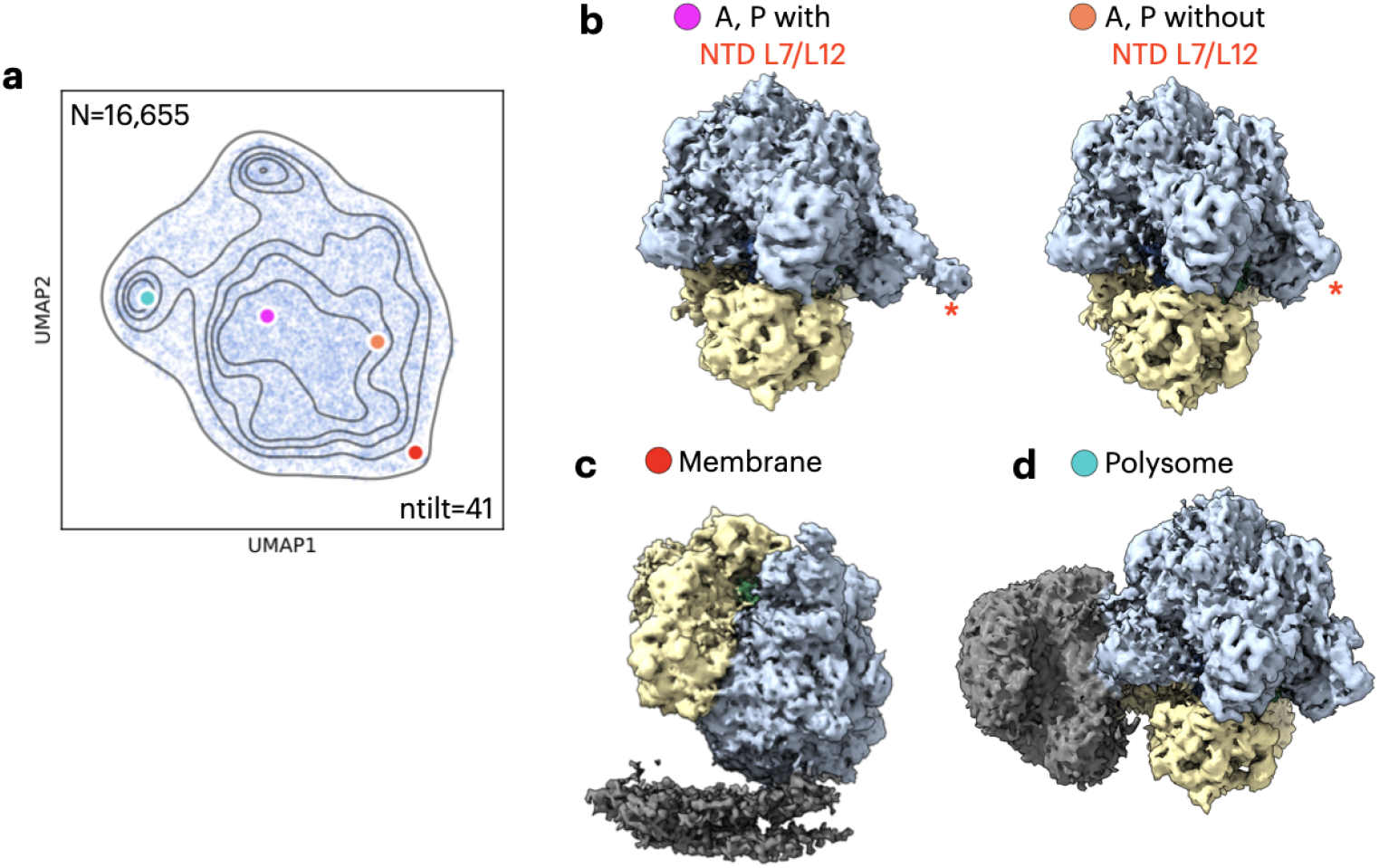
Additional states of the *M. pneumoniae* ribosome identified in cryoDRGN-ET. **a.** UMAP visualization of the latent space from cryoDRGN-ET training (D=128, 3.9 Å/pixel) on the *M. pneumoniae* ribosome filtered particle set, using 41 tilts per particle during training. Latent embeddings for representative density maps are highlighted. **b.** Representative maps with (left) and without (right) density present for the NTD of L7/L12, as highlighted by the red asterisk. **c.** Representative map depicting a membrane-bound ribosome. **d.** Representative map depicting polysome density.

**Fig. S7.**
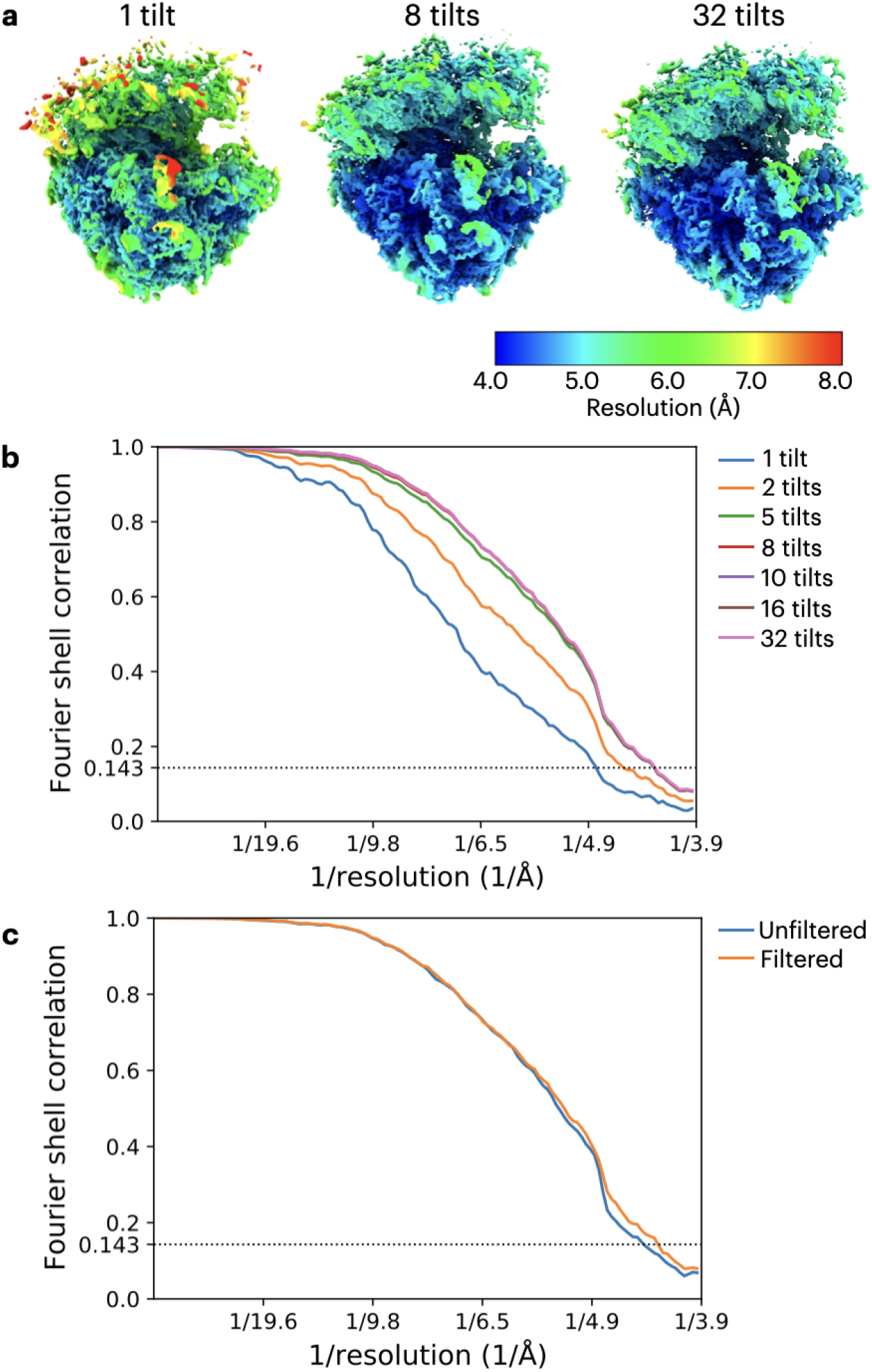
Homogeneous reconstruction of the *S. cerevisiae* ribosome varying the number of tilt images (D=256, 1.96 Å/pixel). **a.** Local resolution estimated from RELION 4.0 [42] for reconstructions using 1 tilt, 8 tilt, and 32 tilts per particle, with maps obtained through voxel-based backprojection in cryoDRGN. **b.** GSFSC curves for varying numbers of tilts per particles. **b.** GSFSC curves for either the full particle set (119,031 particles) or the filtered particle set (93,281 particles) using all tilts per particle.

**Fig. S8.**
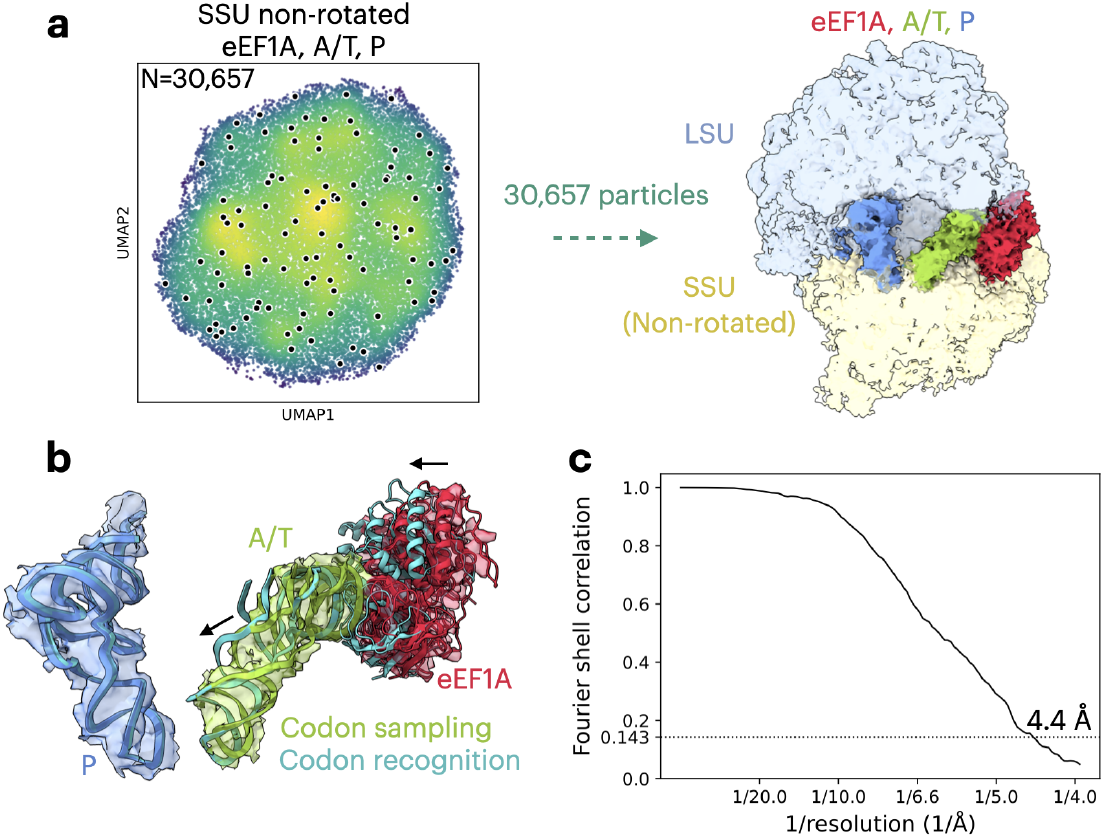
Homogeneous reconstruction of translational states of the *S. cerevisiae* ribosome identified in cryoDRGN-ET in the eEF1A, A/T, P state. **a.** The left panel shows the UMAP visualization of the latent space from cryoDRGN-ET training on the indicated particle set, with overlaid heatmaps highlighting particles belonging to each state. The right column depicts the homogeneous reconstruction (D=256, 1.96 Å/pixel) from cryoDRGN-ET’s voxel-based backprojection for particles selected in each state. Reconstructions are low-pass filtered to the GSFSC resolution and colored by corresponding factors. **b.** Superposition of the P tRNA, A/T tRNA, and eEF1A from this state vs the codon recognition state (cyan) from PDB ID: 5LZS [25]. The P tRNA, A/T tRNA, and eEF1A were separately docked into the reconstruction from **a)** for comparison to the codon recognition state. Density is shown from the reconstruction in **a)** around these factors. Arrows indicate the shift in position of the A/T tRNA and eEF1A between the codon sampling and codon recognition states. **c.** GSFSC curves for the eEF1A, A/T, P state.

**Fig. S9.**
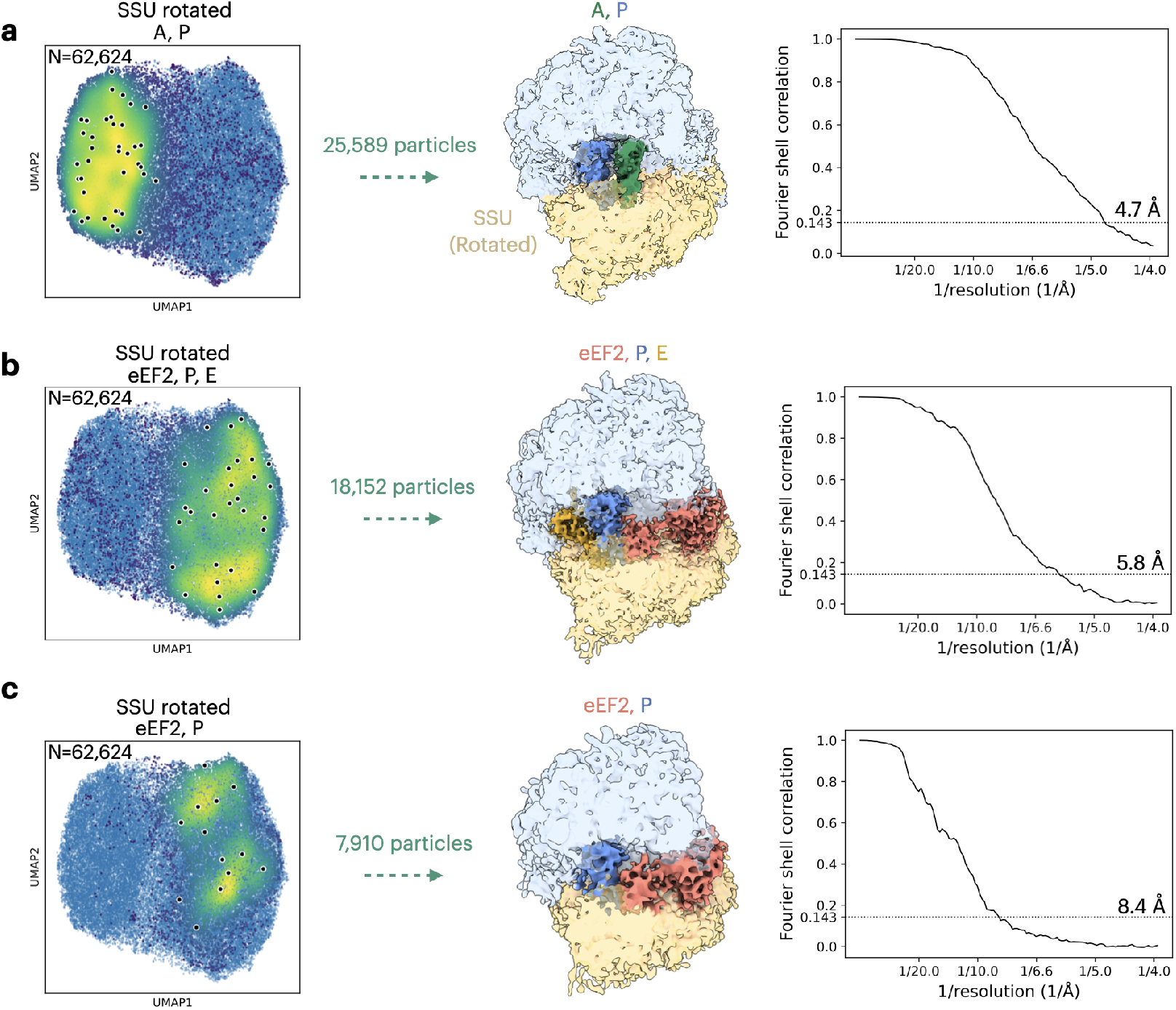
Homogeneous reconstruction of translational states of the *S. cerevisiae* ribosome identified in cryoDRGN-ET in the following states: **a.** the A, P state; **b.** the eEF2, P, E state; and **c.** the eEF2, P state. For each state, the left panel shows the UMAP visualization of the latent space from cryoDRGN-ET training on the indicated particle set, with overlaid heatmaps highlighting particles belonging to each state. The middle column depicts the homogeneous reconstruction (D=256, 1.96 Å/pixel) from cryoDRGN-ET’s voxel-based backprojection for particles selected in each state. Reconstructions are low-pass filtered to the GSFSC resolution and colored by corresponding factors. In the right column, GSFSC curves are depicted for each case.

**Fig. S10.**
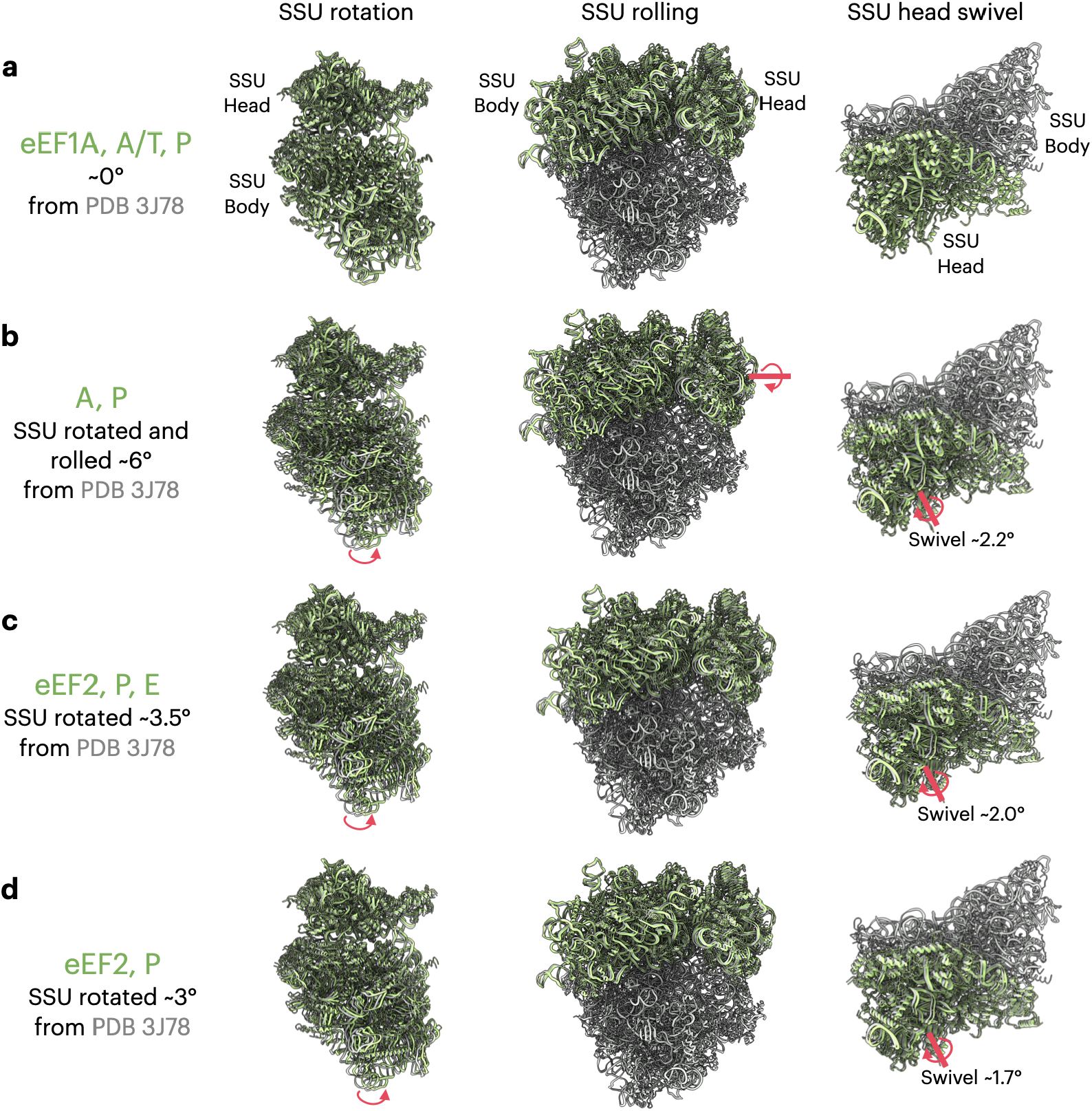
Conformational motions of the *S. cerevisiae* ribosome SSU identified in cryoDRGN in the following states: **a.** the eEF1A, A/T, P state; **b.** the A, P state; **c.** the eEF2, P, E state; and **d.** the eEF2, P state. In all panels, the non-rotated *S. cerevisiae* ribosome structure in PDB ID 3J78 [55] is shown in grey, and docked models into high-resolution reconstructions from cryoDRGN-ET are shown in green. The left column shows the rotation of the SSU, with models aligned on the LSU (LSU not shown for clarity) and red arrows indicating cases of significant rotation. The middle column shows rolling of the SSU, again with models aligned on the LSU and red arrows indicating the case with significant rolling. Finally, the right column shows the SSU head swivel, with models aligned on the SSU body (LSU removed and not shown), and red arrows indicating cases where a minor head swivel is present. Rotations angles between docked models and coordinates from PDB ID 3J78 [55] are measured in ChimeraX [43]. For each state we report two rotation angles. The angle representing either “SSU rotation” or “SSU rolling and rotation” is measured as the rotation angle required to superimpose the state’s SSU onto the SSU of PDB 3J78, when structures aligned on the LSU. The angle representing the head swivel is measured as the rotation angle required to superimpose the SSU head between that state and PDB ID 3J78, when the SSU structures are aligned on the SSU body.

**Fig. S11.**
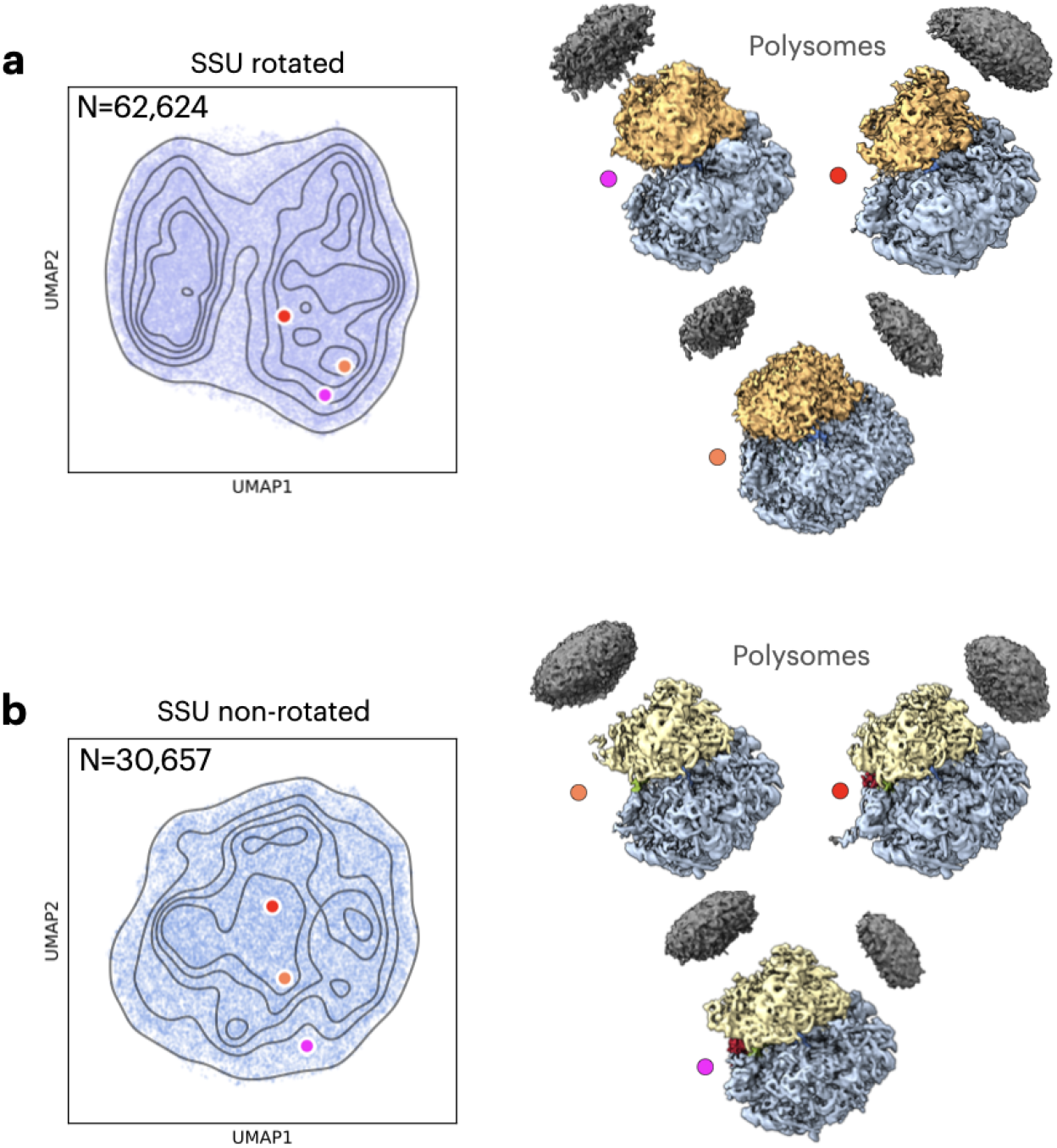
Polysome structures of the *S. cerevisiae* ribosome SSU identified in cryoDRGN-ET from **a.** SSU rotated and **b.** SSU non-rotated particles. In the left column, UMAP visualization of the latent space from cryoDRGN-ET training (D=128, 3.92 Å/pixel) on the indicated particle set. Latent embeddings for representative density maps are highlighted. In the right column, representative density maps from cryoDRGN-ET depicting polysome density are shown.

**Fig. S12.**
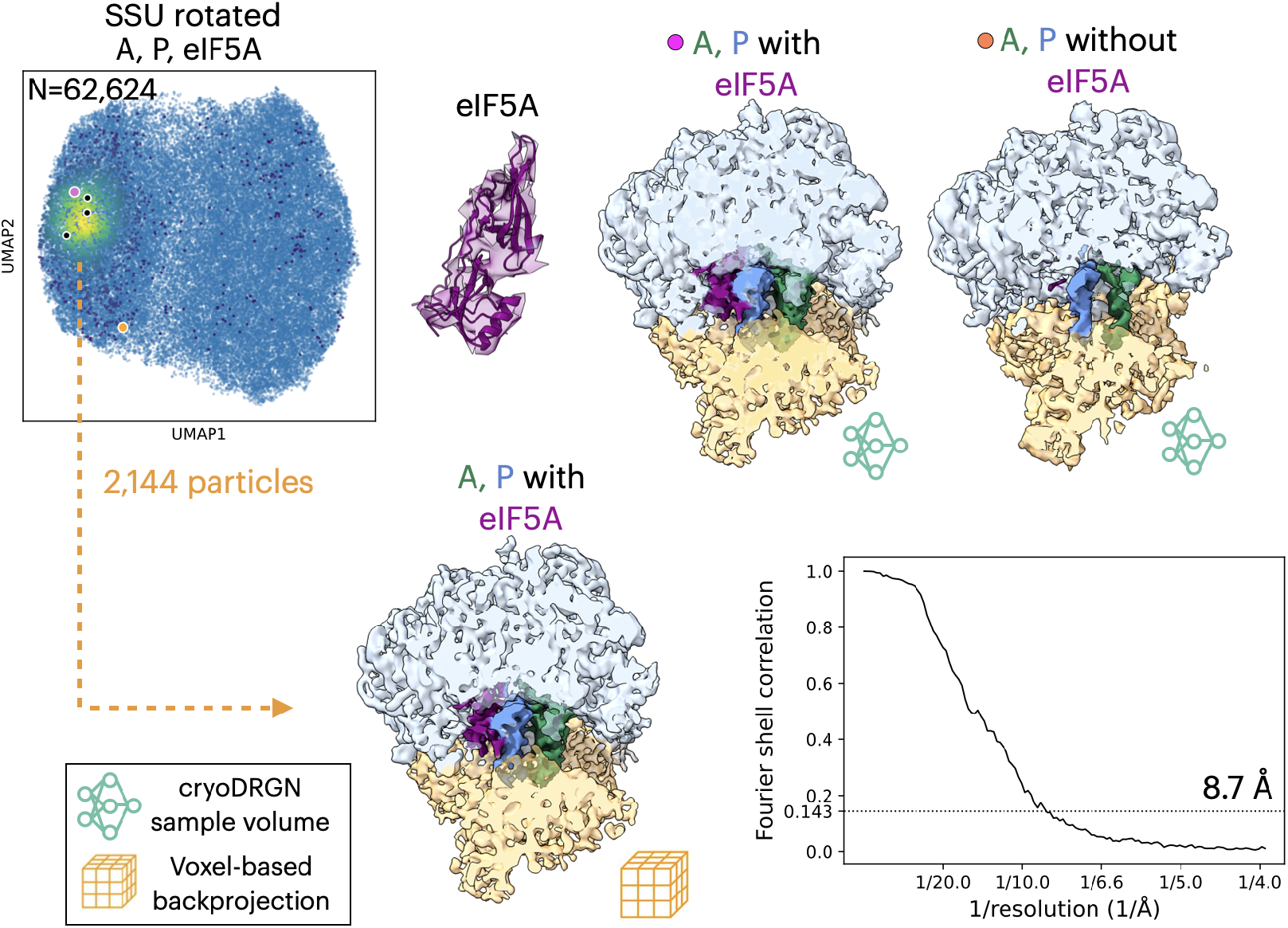
*S. cerevisiae* ribosome structures with the eIF5A initiation factor from cryoDRGN-ET along with validation from voxel-based backprojection. The top left panel shows the UMAP visualization of the latent space from cryoDRGN-ET training on the SSU rotated particle set, with the overlaid heatmap highlighting particle classes with representative density maps including eIF5A density. The top right includes representative maps from cryoDRGN-ET with and without eIF5A density, with their latent embeddings indicated in the UMAP visualization. An atomic model for eIF5A in shown in density from the cryoDRGN-ET representative map (right). On the bottom left, a reconstruction from cryoDRGN-ET’s voxel-based backprojection is shown low-pass filtered to GSFSC resolution, and on the bottom right, the GSFSC curve for this reconstruction is shown.

**Fig. S13.**
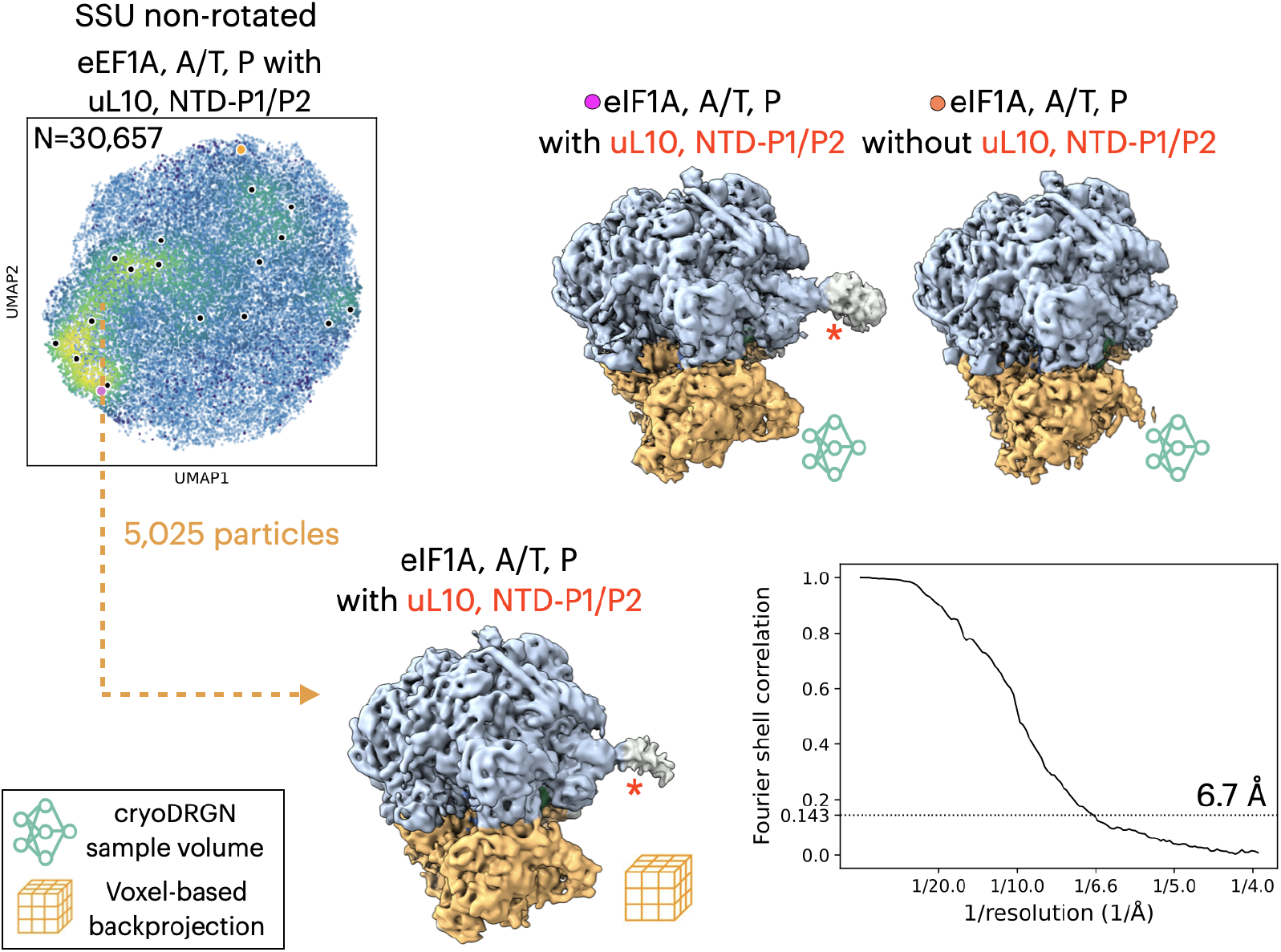
*S. cerevisiae* ribosome structures with variable occupancy of the uL10, NTD-P1/P2 from cryoDRGN-ET along with validation from voxel-based backprojection. The top left panel shows the UMAP visualization of the latent space from cryoDRGN-ET training on the SSU non-rotated particle set, with the overlaid heatmap highlighting particle classes with representative density maps including density for uL10 and the NTD of P1 and P2 heterodimers. The top right includes representative maps from cryoDRGN-ET with and without uL10(P1-P2)_2_ density, with their latent embeddings indicated in the UMAP visualization and the relevant density highlighted with a red asterisk. On the bottom left, a reconstruction from cryoDRGN-ET’s voxel-based backprojection is shown low-pass filtered to GSFSC resolution (0.143 cutoff), and on the bottom right, the GSFSC curve for this reconstruction is shown.

**Fig. S14.**
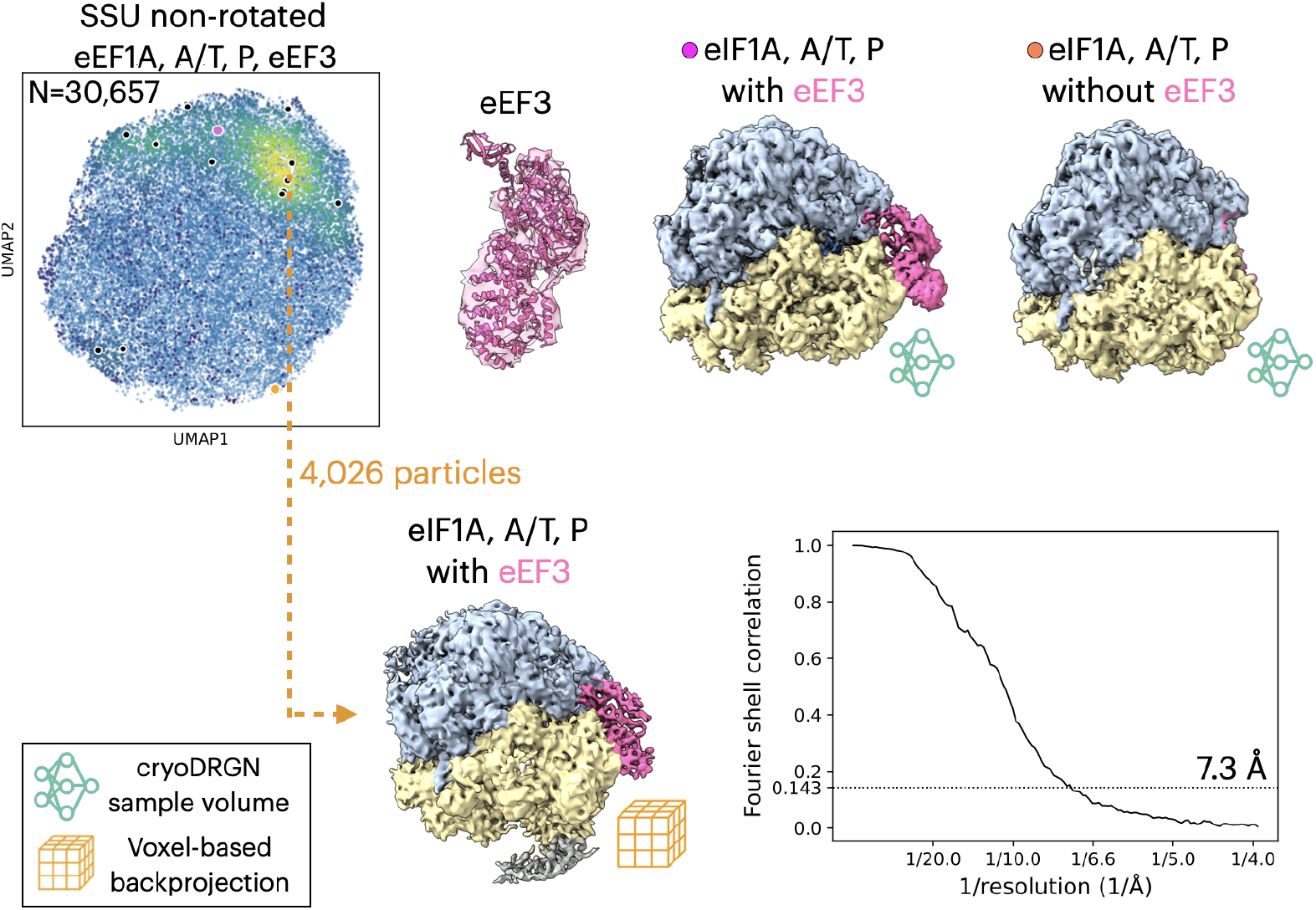
*S. cerevisiae* ribosome structures with eEF3 from cryoDRGN-ET and validated along with validation from voxel-based backprojection. The top left panel shows the UMAP visualization of the latent space from cryoDRGN-ET training on the SSU non-rotated particle set, with the overlaid heatmap highlighting particle classes with representative density maps including eEF3 density. The top right includes representative maps from cryoDRGN-ET with and without eEF3 density, with their latent embeddings indicated in the UMAP visualization. An atomic model for eEF3 in shown in density from the cryoDRGN-ET representative map (right). On the bottom left, a reconstruction from cryoDRGN-ET’s voxel-based backprojection is shown low-pass filtered to GSFSC resolution (0.143 cutoff), and on the bottom right, the GSFSC curve for this reconstruction is shown.

## B SUPPLEMENTARY VIDEOS

**Supplementary Video 1**Representative cryoDRGN-ET density maps of the *M. pneumoniae* 70S ribosome from kmeans clustering (k=100) colored according to translational state as in Fig. 2. 7 unclassified maps are excluded.

**Supplementary Video 2** CryoDRGN-ET continuous trajectory of the *M. pneumoniae* 70S ribosome generated by interpolating through four translational states. 10 density maps are generated along the interpolation path between consecutive states, and maps are colored according to each translational state as in Fig. 2. The interpolation path is shown overlaid on a UMAP visualization of the latent space.

**Supplementary Video 3** CryoDRGN-ET continuous trajectory of the *M. pneumoniae* 70S ribosome generated by interpolating through the latent embeddings of the systematically sampled representative density maps in Supplementary Video 1. The interpolation path is shown overlaid on a UMAP visualization of the latent space.

**Supplementary Video 4** CryoDRGN-ET continuous trajectory of the *S. cerevisiae* 80S ribosome generated by interpolating through the identified translational states. 10 density maps are generated along the interpolation path between consecutive states, and maps are colored according to each translational state as in Fig. 3. The interpolation path is shown overlaid on a UMAP visualization of the latent space.

**Supplementary Video 5**CryoDRGN-ET continuous trajectory of the *S. cerevisiae* 80S ribosome generated by interpolating through the latent embeddings of systematically sampled representative density maps. The interpolation path is shown overlaid on a UMAP visualization of the latent space.

## REFERENCES

[1] Oikonomou, C. M. & Jensen, G. J. Cellular electron cryotomography: Toward structural biology in situ. Annu. Rev. Biochem. 86, 873–896 (2017).

[2] Galaz-Montoya, J. G. & Ludtke, S. J. The advent of structural biology in situ by single particle cryo-electron tomography. Biophys Rep 3, 17–35 (2017).

[3] Tegunov, D., Xue, L., Dienemann, C., Cramer, P. & Mahamid, J. Multi-particle cryo-EM refinement with M visualizes ribosome-antibiotic complex at 3.5 å in cells. Nature Methods 18, 186–193 (2021).

[4] Zivanov, J. et al. A Bayesian approach to single-particle electron cryo-tomography in RELION-4.0. Elife 11 (2022).

[5] Himes, B. A. & Zhang, P. emClarity: software for high-resolution cryo-electron tomography and subtomogram averaging. Nat. Methods 15, 955–961 (2018).

[6] Chen, M. et al. A complete data processing workflow for cryo-ET and subtomogram averaging. Nat. Methods 16, 1161–1168 (2019).

[7] Wan, W., Khavnekar, S., Wagner, J., Erdmann, P. & Baumeister, W. STOPGAP: a software package for subtomogram averaging and refinement. Microscopy and Microanalysis 26, 2516–2516 (2020).

[8] Khavnekar, S., et al. Optimizing Cryo-FIB lamellas for sub-5Å in situ structural biology (2022).

[9] Harastani, M., Eltsov, M., Leforestier, A. & Jonic, S. HEMNMA-3D: Cryo electron tomography method based on normal mode analysis to study continuous conformational variability of macromolecular complexes. Front Mol Biosci 8, 663121 (2021).

[10] Harastani, M., Eltsov, M., Leforestier, A. & Jonic, S. TomoFlow: Analysis of continuous conformational variability of macromolecules in cryogenic subtomograms based on 3D dense optical flow. J. Mol. Biol. 434, 167381 (2022).

[11] Erdmann, P. S. et al. In situ cryo-electron tomography reveals gradient organization of ribosome biogenesis in intact nucleoli. Nature Communications 12, 5364 (2021).

[12] Xue, L. et al. Visualizing translation dynamics at atomic detail inside a bacterial cell. Nature 610, 205–211 (2022).

[13] Hoffmann, P. C. et al. Structures of the eukaryotic ribosome and its translational states in situ. Nat. Commun. 13, 7435 (2022).

[14] Xing, H. et al. Translation dynamics in human cells visualized at high resolution reveal cancer drug action. Science 381, 70–75 (2023).

[15] Zhong, E. D., Bepler, T., Berger, B. & Davis, J. H. CryoDRGN: reconstruction of heterogeneous cryo-EM structures using neural networks. Nature methods 18, 176–185 (2021).

[16] Punjani, A. & Fleet, D. J. 3DFlex: determining structure and motion of flexible proteins from cryo-EM. Nature Methods 1–11 (2023).

[17] Chen, M. & Ludtke, S. J. Deep learning-based mixed-dimensional gaussian mixture model for characterizing variability in cryo-EM. Nature methods 18, 930–936 (2021).

[18] Xie, Y., et al. Neural fields in visual computing and beyond (2021). 2111.11426.

[19] Grant, T. & Grigorieff, N. Measuring the optimal exposure for single particle cryo-EM using a 2.6 å reconstruction of rotavirus VP6. Elife 4, e06980 (2015).

[20] McInnes, L., Healy, J. & Melville, J. UMAP: Uniform manifold approximation and projection for dimension reduction (2018). 1802.03426.

[21] Montesano-Roditis, L., Glitz, D. G., Traut, R. R. & Stewart, P. L. Cryo-electron microscopic localization of protein L7/L12 within the escherichia coli 70 S ribosome by difference mapping and nanogold labeling. Journal of Biological Chemistry 276, 14117–14123 (2001).

[22] Kater, L. et al. Partially inserted nascent chain unzips the lateral gate of the Sec translocon. EMBO reports 20, e48191 (2019).

[23] Brandt, F. et al. The native 3D organization of bacterial polysomes. Cell 136, 261–271 (2009).

[24] Powell, B. M. & Davis, J. H. Learning structural heterogeneity from cryo-electron sub-tomograms with tomoDRGN (2023).

[25] Shao, S. et al. Decoding mammalian ribosome-mRNA states by translational GTPase complexes. Cell 167, 1229–1240 (2016).

[26] Choi, A. K., Wong, E. C., Lee, K.-M. & Wong, K.-B. Structures of eukaryotic ribosomal stalk proteins and its complex with trichosanthin, and their implications in recruiting ribosome-inactivating proteins to the ribosomes. Toxins 7, 638–647 (2015).

[27] Ranjan, N. et al. Yeast translation elongation factor eEF3 promotes late stages of tRNA translocation. The EMBO journal 40, e106449 (2021).

[28] Zhong, E. D., Lerer, A., Davis, J. H. & Berger, B. CryoDRGN2: Ab initio neural reconstruction of 3D protein structures from real cryo-EM images. In Proceedings of the IEEE/CVF International Conference on Computer Vision, 4066–4075 (2021).

[29] Levy, A., Wetzstein, G., Martel, J. N., Poitevin, F. & Zhong, E. Amortized inference for heterogeneous reconstruction in cryo-EM. Advances in Neural Information Processing Systems 35, 13038–13049 (2022).

[30] Tancik, M., et al. Fourier features let networks learn high frequency functions in low dimensional domains (2020). 2006.10739.

[31] Zhong, E. D., Bepler, T., Davis, J. H. & Berger, B. Reconstructing continuous distributions of 3d protein structure from cryo-EM images. In International Conference on Learning Representations (ICLR) (2020).

[32] Zhong, E. et al. zhonge/cryodrgn: Version 1.0.0-beta (2022). URL https://doi.org/10.5281/zenodo.6554048.

[33] Grant, T. & Grigorieff, N. Measuring the optimal exposure for single particle cryo-em using a 2.6 å reconstruction of rotavirus VP6. elife 4, e06980 (2015).

[34] Bharat, T. A., Russo, C. J., Löwe, J., Passmore, L. A. & Scheres, S. H. Advances in single-particle electron cryomicroscopy structure determination applied to sub-tomogram averaging. Structure 23, 1743–1753 (2015).

[35] Hagen, W. J., Wan, W. & Briggs, J. A. Implementation of a cryo-electron tomography tilt-scheme optimized for high resolution subtomogram averaging. Journal of structural biology 197, 191–198 (2017).

[36] Kingma, D. P. & Welling, M. Auto-encoding variational bayes (2022). 1312.6114.

[37] Kingma, D. P. & Ba, J. Adam: A method for stochastic optimization (2014). 1412.6980.

[38] Sindelar, C. V. & Grigorieff, N. Optimal noise reduction in 3d reconstructions of single particles using a volume-normalized filter. Journal of structural biology 180, 26–38 (2012).

[39] Rosenthal, P. B. & Henderson, R. Optimal determination of particle orientation, absolute hand, and contrast loss in single-particle electron cryomicroscopy. J. Mol. Biol. 333, 721–745 (2003).

[40] Zheng, S., et al. AreTomo: An integrated software package for automated marker-free, motion-corrected cryo-electron tomographic alignment and reconstruction. Journal of Structural Biology: X 6, 100068 (2022).

[41] Wagner, T. et al. SPHIRE-crYOLO is a fast and accurate fully automated particle picker for cryo-EM. Commun Biol 2, 218 (2019).

[42] Kimanius, D., Dong, L., Sharov, G., Nakane, T. & Scheres, S. H. W. New tools for automated cryo-EM single-particle analysis in RELION-4.0. Biochem. J 478, 4169–4185 (2021).

[43] Pettersen, E. F. et al. UCSF ChimeraX: Structure visualization for researchers, educators, and developers. Protein Science 30, 70–82 (2021).

[44] Khavnekar, S., Erdmann, P. & Wan, W. TOMOMAN: Streamlining cryo-electron tomography and subtomogram averaging workflows using TOMOgram MANager (2023).

[45] Bai, X.-c., Fernandez, I. S., McMullan, G. & Scheres, S. H. Ribosome structures to near-atomic resolution from thirty thousand cryo-EM particles. elife 2, e00461 (2013).

[46] Rohou, A. & Grigorieff, N. CTFFIND4: Fast and accurate defocus estimation from electron micrographs. Journal of structural biology 192, 216–221 (2015).

[47] Mastronarde, D. N. & Held, S. R. Automated tilt series alignment and tomographic reconstruction in IMOD. Journal of structural biology 197, 102–113 (2017).

[48] Turoňová, B., Schur, F. K., Wan, W. & Briggs, J. A. Efficient 3D-CTF correction for cryo-electron tomography using NovaCTF improves subtomogram averaging resolution to 3.4 å. Journal of structural biology 199, 187–195 (2017).

[49] Pellegrino, S. et al. Structural insights into the role of diphthamide on elongation factor 2 in mRNA reading-frame maintenance. Journal of molecular biology 430, 2677–2687 (2018).

[50] Himes, B. & Grigorieff, N. Cryo-TEM simulations of amorphous radiation-sensitive samples using multislice wave propagation. IUCrJ 8, 943–953 (2021).

[51] Tegunov, D. & Cramer, P. Real-time cryo-electron microscopy data preprocessing with Warp. Nature methods 16, 1146–1152 (2019).

[52] Zivanov, J. et al. New tools for automated high-resolution cryo-em structure determination in RELION-3. elife 7, e42166 (2018).

[53] Voorhees, R. M., Fernández, I. S., Scheres, S. H. & Hegde, R. S. Structure of the mammalian ribosome-Sec61 complex to 3.4 å resolution. Cell 157, 1632–1643 (2014).

[54] Buschauer, R. et al. The Ccr4-Not complex monitors the translating ribosome for codon optimality. Science 368, eaay6912 (2020).

[55] Svidritskiy, E., Brilot, A. F., San Koh, C., Grigorieff, N. & Korostelev, A. A. Structures of yeast 80S ribosome-tRNA complexes in the rotated and nonrotated conformations. Structure 22, 1210–1218 (2014).

